# Π-Π Interactions Stabilize PeptoMicelle-Based Formulations of Pretomanid Derivatives Leading to Promising Therapy Against Tuberculosis in Zebrafish and Mouse Models

**DOI:** 10.1101/2022.03.10.483770

**Authors:** Nils-Jørgen K. Dal, Gabriela Schäfer, Andrew M. Thompson, Sascha Schmitt, Natalja Redinger, Noelia Alonso-Rodriguez, Kerstin Johann, Jessica Ojong, Jens Wohlmann, Andreas Best, Kaloian Koynov, Rudolf Zentel, Ulrich E. Schaible, Gareth Griffiths, Matthias Barz, Federico Fenaroli

## Abstract

Tuberculosis is the deadliest bacterial disease globally, threatening the lives of millions every year. New antibiotic therapies that can shorten the duration of treatment, improve cure rates, and impede the development of drug resistance are desperately needed. Here, we used polymeric micelles to encapsulate four second-generation derivatives of the antitubercular drug pretomanid that had previously displayed much better *in vivo* activity against *Mycobacterium tuberculosis* than pretomanid itself. Because these compounds were relatively hydrophobic, we expected that such micellar formulations would increase drug bioavailability, reduce toxicities, and improve therapeutic outcomes. The polymeric micelles were based on polypept(o)ides (PeptoMicelles) and were stabilized in their hydrophobic core by π-π interactions, allowing the efficient encapsulation of aromatic pretomanid derivatives. The stability of these π-π-stabilized PeptoMicelles was demonstrated in water, blood plasma, and lung surfactant by fluorescence cross-correlation spectroscopy and was further supported by prolonged circulation times of several days in the vasculature of zebrafish larvae. The pretomanid derivative with the best *in vitro* potency against *Mycobacterium marinum* (“drug D”) was also the most efficacious PeptoMicelle formulation tested in the zebrafish larvae infection model, almost completely eradicating the bacteria at non-toxic doses. This lead formulation was further assessed against *Mycobacterium tuberculosis* in the susceptible C3HeB/FeJ mouse model, which develops human-like necrotic granulomas. Following intravenous administration, the drug D micellar formulation significantly reduced bacterial burden and inflammatory responses in the lungs and spleens of infected mice.

## INTRODUCTION

Tuberculosis (TB), arising from infection by bacteria of the *Mycobacterium tuberculosis* (*Mtb*) complex, remains a major global health issue, with about 10 million new cases and approximately 1.5 million deaths caused by TB in 2020.^1^ Current 6-month treatment for drug-sensitive (DS) TB involves combinations of four first-line antibiotics, isoniazid, rifampicin, pyrazinamide, and ethambutol. These drugs can cause hepatotoxicity, neurotoxicity, and other adverse effects, a common reason for low treatment compliance in some TB patients. Inadequate treatment then promotes the selection of multidrug-resistant (MDR) and even extensively drug-resistant (XDR) *Mtb* strains that are increasingly difficult to eradicate.^2^ While three new drugs, bedaquiline (BDQ), delamanid, and pretomanid, were approved for the treatment of MDR TB in the past decade, more effective therapies are urgently needed.^3^ Notably, soon after the introduction of BDQ and delamanid, *Mtb* isolates resistant to these agents were observed.^4^

Pretomanid is a very promising drug that is currently in phase III combination trials for the treatment of DS and MDR/XDR TB.^3^ Due to its impressive clinical results, pretomanid received early approval in 2019 for use against XDR TB and treatment-intolerant/non-responsive MDR TB, in combination with BDQ and linezolid (BPaL).^5^ Recently, many second-generation analogues of pretomanid have been reported that demonstrate superior potencies *in vitro* against *Mtb* and greatly enhanced therapeutic efficacies in both acute and chronic infection BALB/c mouse models.^6–8^

Importantly, pretomanid and its analogues are active against both replicating *Mtb* and the non-replicating bacteria considered to be responsible for latent TB.^9,10^ Pretomanid kills replicating *Mtb* by inhibiting cell wall mycolate synthesis.^11^ In contrast, nitric oxide release (leading to respiratory poisoning) is responsible for killing the non-replicating bacteria.^10,12^

Antibiotics showing good therapeutic properties *in vitro* are often limited by their hydrophobicity and low bioavailability.^13^ Therefore, nanoparticle (NP) based drug delivery systems have attracted increasing attention for TB treatment.^14–20^ This approach can substantially improve drug bioavailability, facilitate drug administration, and protect both the cargo (*e.g.,* from premature degradation) and the patient (*e.g.*, from drug accumulation and cytotoxicity in healthy tissues).^15,19,21^ Moreover, rapid depletion of drug plasma levels due to excretion or first-pass metabolism can be avoided, and sustained drug release from NPs could enable a reduced frequency of administration.^16,17^ Therefore, in TB therapy, NPs have great potential to increase treatment efficacy and patient compliance,^20,22–25^ which could impede the development of drug resistance.

Among NPs, polymeric micelles (PM) are especially interesting as nano-sized drug delivery systems for hydrophobic drugs, leading to several clinical trials and one approved medicine (Genexol-PM).^13,26–31^ PM can entrap hydrophobic drugs in the micelle core that is shielded from the surrounding medium by a hydrophilic outer shell, facilitating much greater water solubility and other advantages, as described above.^32^ The most prominent hypothesis explaining how NPs can leave the bloodstream involves local inflammation near tumors or TB granulomas; the endothelial lining of the blood vessels becomes leaky, allowing NPs to diffuse out [enhanced permeability and retention (EPR) effect].^33,34^ To maximize this passive accumulation near the infection site, NPs should circulate for extended periods in the bloodstream.^35–37^ Therefore, various physical and chemical strategies for the stabilization of PM and prolongation of drug entrapment have been developed, which include chemical crosslinking or physical crosslinking by non-covalent interactions, *e.g.*, π-π-interactions.^13,27,36,38,39^

Here, we have developed novel PM based on the polypept(o)ide poly(*γ*-benzyl-L-glutamic acid)-*block*-polysarcosine for the delivery of anti-TB drugs.^40^ This amphiphilic polymer combines the stealth-like properties of the hydrophilic polypeptoid polysarcosine (pSar)^41^ as the surface shell with the hydrophobic core polypeptide poly(*γ*-benzyl-L-glutamic acid) (pGlu(OBn)), which contains aromatic benzyl groups in the side chains.^42^ In aqueous solution, these polypept(o)ides form micelles, which we show here to be stabilized by non-covalent π-π interactions between benzyl groups in the core. The attractive potential between aromatic systems is particularly strong when electron-rich and electron-deficient aromatic molecules interact as both π-π and polar interactions (permanent dipoles or even charge transfers) can contribute to the overall stability.^43^ Because the pretomanid derivatives contain electron-deficient aromatic systems, we hypothesized that attractive interactions with electron-rich benzyl groups in the polypeptide block would occur, leading to PM with enhanced stability. Although earlier noted by the group of Kataoka^44^, the concept of micelles stabilized by π-π interactions facilitating better drug delivery was first demonstrated by Hennink and coworkers in 2013 and applied to tumor therapy in 2015.^45,46^

We recently showed that polypept(o)ide-based NPs accumulate in granulomas *via* an EPR-like mechanism in zebrafish larvae and mice.^47^ Zebrafish have a unique advantage over all mammalian organisms in that the embryo and early larvae are transparent, allowing real-time live imaging with high resolution.^48,49^ Zebrafish infected with the bacterium that causes TB in fish and frogs, *Mycobacterium marinum* (*Mm*), have been widely studied as a model system for mammalian TB.^50–52^ *Mm* is closely related to *Mtb* but, whereas *Mtb* grows at 37 °C, *Mm* is a pathogen of cold-blooded animals and grows optimally at around 28 °C, as does the zebrafish. The zebrafish larvae model of TB has been developed into a rapid screening platform to determine (free) drug efficacy.^53,54^ Zebrafish are also increasingly being used for the toxicological screening of compounds.^55,56^

The current study has investigated the utility of π-π-PeptoMicelles as a delivery system for pretomanid and four second-generation derivatives. Following *in vitro* testing of the free drugs and their encapsulation in PM, the resulting formulations were tested for toxicity and therapeutic efficacy in zebrafish larvae infected with *Mm*. From these data, we selected the most efficacious formulation for further evaluation against *Mtb* in the susceptible C3HeB/FeJ mouse model of TB.^57^

## RESULTS AND DISCUSSION

### *In vitro* antimycobacterial activity and cytotoxicity in VERO cells

Pretomanid (PA-824, **1**; **Figure 1**) and four second-generation analogues (drugs A-D) were initially screened *in vitro* against liquid cultures of *Mtb* (strain H37Rv) and *Mm* (strain M). Newly measured MIC data against *Mtb* (**Table 1**) were all within experimental errors of published values.^6–8^ Concerning *Mm*, pretomanid itself was poorly active (MIC_90_ 209 µM), broadly in line with results reported by Lee et al. (MIC 45 µM)^58^ and Dalton et al. (MIC >10 µM).^59^ (Contrary to one report,^59^ pretomanid also failed to show any *in vivo* activity against *Mm* using the zebrafish larvae model described below; **Figure S11**). Less soluble analogues A-D exhibited better activity against *Mm in vitro*, with drug D being the most effective (MIC_90_ 0.75 µM). These compounds were further examined for cytotoxicity toward mammalian cells (VERO) in a 72 h cell viability assay, but none displayed a significant liability (IC_50_s >200 µM).

**Figure 1.**
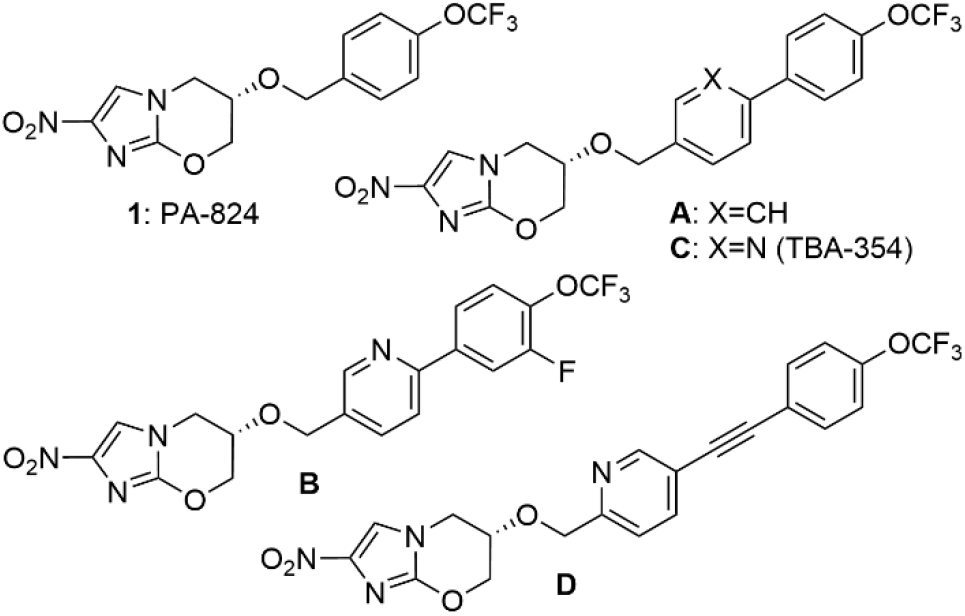
Chemical structures of pretomanid (**1**) and four efficacious derivatives.

**Table 1.** Solubility, inhibitory potency, and cytotoxicity data for pretomanid and four derivatives.

| Compound | Solubility<sup>a</sup><br>($\mu\text{g/mL}$) | MIC (*Mtb*)<sup>b</sup><br>($\mu\text{M}$) | MIC<sub>90</sub><br>(*Mm*)<sup>b</sup> ($\mu\text{M}$) | IC<sub>50</sub> (VERO)<br>($\mu\text{M}$) | Selectivity Index<sup>c</sup> | |
| --- | --- | --- | --- | --- | --- | --- |
|  |  |  |  |  | *Mtb* | *Mm* |
| Pretomanid | 19 | 0.51 ± 0.31 | 209 ± 70 | >200 | >392 | >0.95 |
| Drug A | 1.2 | 0.029 ± 0.019 | 1.3 ± 0.6 | >200 | >6896 | >153 |
| Drug B | 3.0 | 0.028 ± 0.018 | 6.4 ± 2.0 | >200 | >7142 | >31 |
| Drug C | 2.3 | 0.025 ± 0.017 | 7.6 ± 4.8 | >200 | >8000 | >26 |
| Drug D | 0.36 | 0.015 ± 0.009 | 0.75 ± 0.40 | >200 | >13333 | >266 |

<sup>a</sup>Solubility in water at 20 °C; data from refs 6 and 7. <sup>b</sup>Minimum inhibitory concentrations against *Mtb* or *Mm* by the REMA assay (see Methods); values are the mean of at least three independent determinations. <sup>c</sup>Selectivity Index (SI), defined by the ratio of VERO IC<sub>50</sub> to MIC (SI = IC<sub>50</sub>/MIC).

### Synthesis and characterization of polypept(o)ides and polymeric micelles for drug encapsulation

The amphiphilic block copolypept(o)ide pGlu(OBn)_27_-*b*-pSar_182_, required for the preparation of PM (π-π-PeptoMicelles), was made by ring-opening polymerization of the appropriate *N*-carboxyanhydrides (NCAs), as described by Birke et al.^42^ (**Scheme 1A**). Characterization by ^1^H NMR spectroscopy, diffusion-ordered spectroscopy (DOSY), and gel permeation chromatography (GPC) confirmed the successful synthesis of block copolymers (**Figures S1-S3**) with narrow molecular weight distributions (Ð<1.2). We chose this polypept(o)ide for the preparation of PM for several reasons. First, pSar provides “stealth” properties for the shielding of NPs in physiological environments and avoids protein corona formation.^60,61^ Moreover, while pSar shows identical solution properties to PEG (solubility and main-chain flexibility in aqueous solution),^61^ it displays an improved immunogenicity^62,63^ and toxicity profile.^64^ Furthermore, as pSar is based on the endogenous amino acid sarcosine, its degradation products can be expected to possess high biocompatibility.^65^ The other component of our NPs, pGlu(OBn), is a hydrophobic polypeptide containing electron-rich benzyl groups in the amino acid side chain. Therefore, we can expect stabilization of the drug formulations due to π-π-interactions between the aromatic groups of pGlu(OBn) and the electron-poor aromatic systems of the synthesized pretomanid derivatives.^36^ However, the overall block length was critical, as shorter pGlu(OBn) segments failed to stabilize the drugs efficiently, while larger hydrophobic blocks resulted in significantly lower drug loading (<10 wt%) and very polydisperse micelles.

**Scheme 1.**
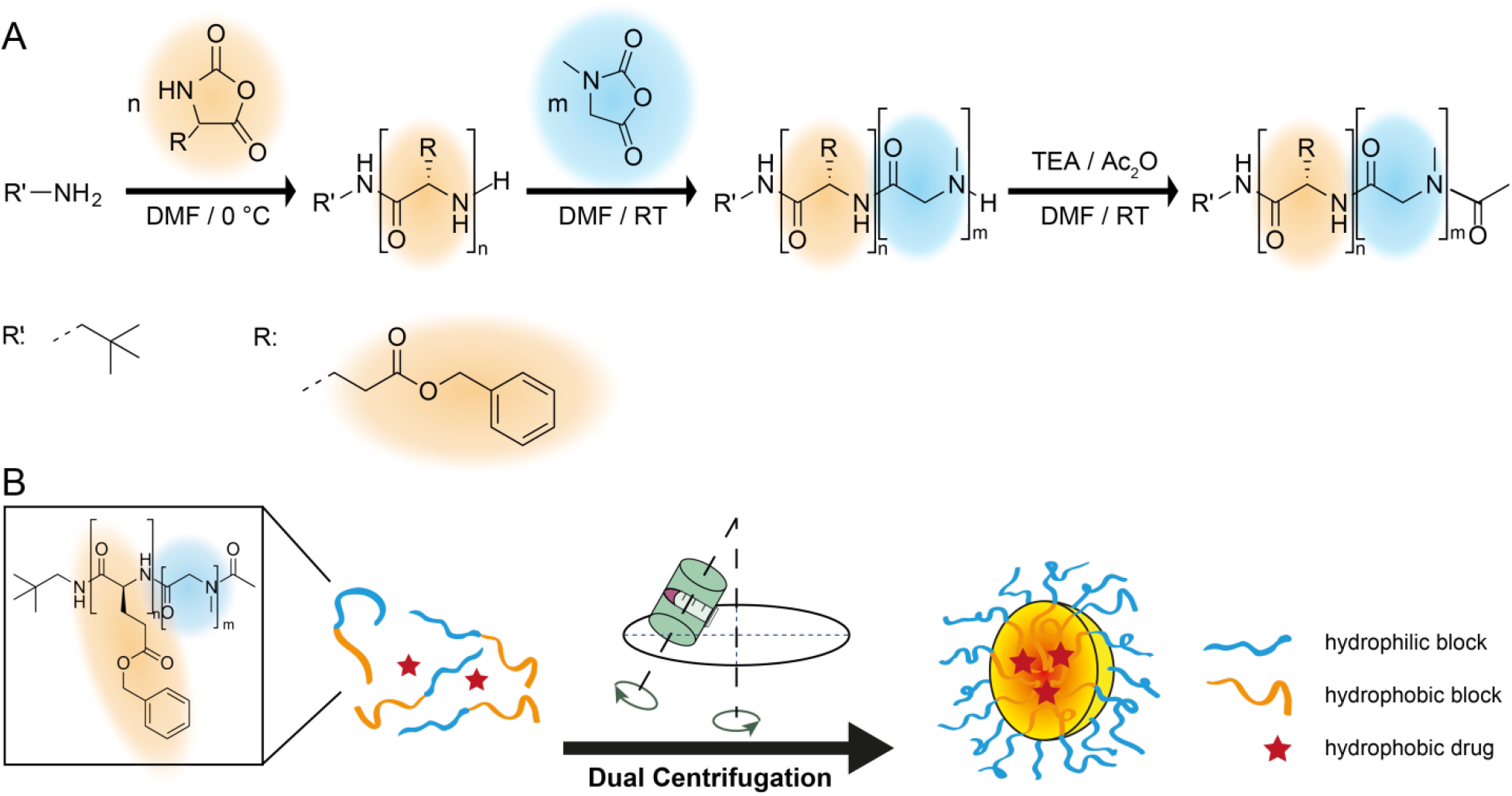
**A:** Synthesis of amphiphilic copolypept(o)ides (pGlu(OBn)-*b*-pSar) by ring-opening NCA polymerization. **B**: Schematic representation of micelle preparation by dual centrifugation (DC).

In contrast to former work from our group, the PeptoMicelles were prepared from the synthesized amphiphilic block copolypept(o)ide by dual centrifugation^66^ (also known as dual asymmetric centrifugation; **Scheme 1B**). In this method, high shear forces are created by the rotation of a viscous sample around its own vertical axis and rotation around the main axis of the centrifuge, thereby enabling more efficient sample homogenization. Furthermore, samples can easily be prepared under reproducible and sterile conditions, which is particularly important for NP samples intended for *in vitro* or *in vivo* testing, long-term storage, and eventually, commercialization.

To obtain micellar formulations of pretomanid and four different hydrophobic derivatives (drugs A-D; **Figure 1**), 30 wt% of the drug was added to the block copolypept(o)ide, and the mixture was subjected to dual centrifugation, as previously reported for non-loaded PeptoMicelles.^47^ The prepared formulations were subsequently characterized by single-angle dynamic light scattering (DLS). The formulations containing pretomanid or the derivatives B, C, and D displayed an average hydrodynamic size comparable to that of the non-loaded (drug-free) PM (*D*_h_ = 84-100 nm; **Table 2**), with acceptable polydispersity indices (≤0.15). Only the formulation containing the more hydrophobic drug A displayed a significantly higher average size (*D*_h_ = 132 nm). Also, the intensity-weighed size distributions of non-loaded PeptoMicelles and the micellar formulation containing drug D (which was later identified as the lead formulation for *in vivo* studies; see below) are shown in **Figure 2A**.

**Figure 2.**
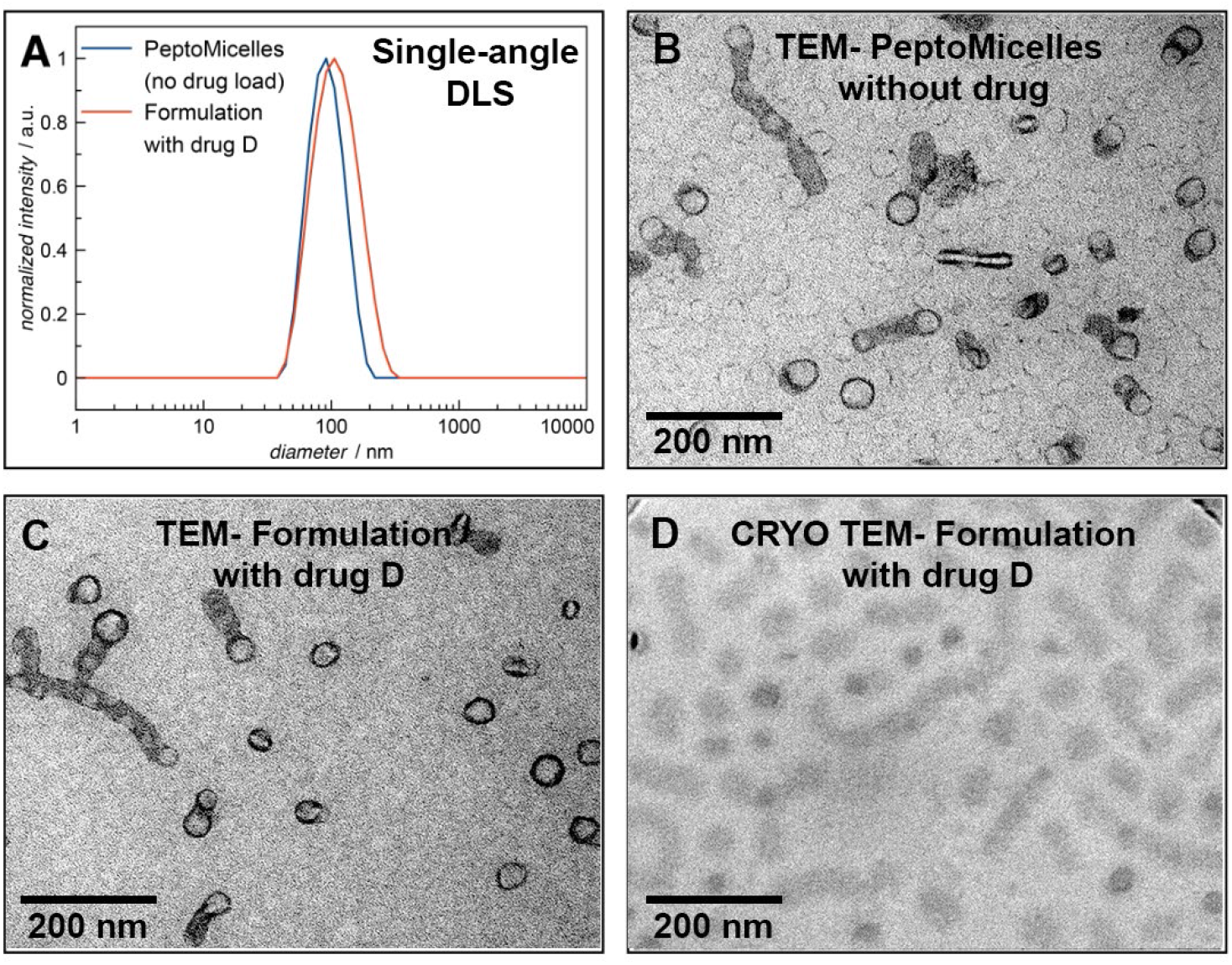
**A:** Characterization of drug-free PeptoMicelles and the micellar drug D formulation by single-angle DLS. **B** and **C:** TEM of PeptoMicelles without or with drug D, respectively. **D:** Cryo TEM.

**Table 2.** Overview of PeptoMicelles and drug-loaded micellar formulations prepared by dual centrifugation.

| Cargo | $D_h^a$ / nm<br>(DLS) | PDI<sup>b</sup> | $D_h$ / nm<br>(FCS) | Zeta potential<br>(mV) | Encapsulation<br>efficiency<sup>c</sup> (%) |
| --- | --- | --- | --- | --- | --- |
| - | 87 ± 1 | 0.09 ± 0.01 | 91 ± 9 | -3.7 ± 0.1 | - |
| Pretomanid | 98 ± 1 | 0.15 ± 0.01 | 102 ± 10 | -3.0 ± 0.2 | 97.1 |
| Drug A | 132 ± 2 | 0.21 ± 0.01 | 126 ± 12 | - 1.9 ± 0.3 | 96.8 |
| Drug B | 84 ± 1 | 0.11 ± 0.01 | 88 ± 8 | -1.8 ± 0.6 | 95.4 |
| Drug C | 90 ± 1 | 0.11 ± 0.05 | 92 ± 10 | -2.5 ± 0.4 | 94.8 |
| Drug D | 100 ± 1 | 0.14 ± 0.01 | 98 ± 10 | -2.7 ± 0.6 | 97.2 |

<sup>a</sup>Z-average sizes were determined by single-angle dynamic light scattering (DLS) at a scattering angle of 173°.
<sup>b</sup>Polydispersity index. All values signify mean ± standard error of the mean (S.E.M.) from $n = 3$ . <sup>c</sup>Encapsulation efficiency was calculated using the equation: $EE\% = \frac{m_{total} - m_{free}}{m_{total}} \times 100\%$

Further measurements demonstrated that all drug-loaded and non-loaded PeptoMicelles possess a neutral zeta (ξ) potential (**Table 2**). The drug encapsulation efficiencies were also excellent (94.8 to 97.2%, **Figure S6**), as we expected for poorly water-soluble molecules. Moreover, drug release kinetic data for these five drug-loaded PM (in PBS buffer at 37 °C) indicated that the micellar formulations were highly stable (**Figure S7**). For PM containing pretomanid derivatives A-D, minimal drug quantities (1-6%) were released over 192 h (the drug D formulation being the most stable). Only for the pretomanid-containing micelles did we encounter higher drug release (∼16% after 192 h).

Non-loaded PeptoMicelles and the micellar formulation of drug D were further characterized by transmission electron microscopy (TEM), using negative staining. TEM images revealed the coexistence of spherical and worm-like micelles for both samples (**Figures 2B** and **2C**). While the diameter of the spherical particles and width of the worm-like micelles were generally below 40 nm, the length of the tubes was found to vary. An additional analysis of the drug D-containing formulation by cryo TEM (**Figure 2D**) displayed the same morphology observed by negative staining TEM. Importantly, similar size distributions and morphologies were observed for both non-loaded and drug D-loaded PeptoMicelles, arguing that they are likely to exhibit the same characteristics with respect to circulation times and biodistribution *in vivo*.

### Exchange dynamics of PeptoMicelles

For a successful *in vivo* application, the stability of the developed micellar formulations under physiological conditions is crucial. If drug-loaded PM disassemble quickly upon injection into the bloodstream, they solubilize the drug but do not change its pharmacokinetic profile.^13^ Nevertheless, an increase in bioavailability without the need for toxic solubilizers, such as Cremophor®, is still a significant therapeutic benefit.^21^ Non-cross-linked PM are usually expected to disassemble rather quickly upon injection due to dilution below their critical micelle concentration (CMC) or binding of the micelle-forming unimers to blood components or cells, which shifts the unimer micelle equilibrium towards the unimer.^13,36^ Therefore, we used fluorescence correlation spectroscopy (FCS)^67^ to investigate the stability of the synthesized PeptoMicelles, with and without pretomanid and drugs A-D. In FCS, fluorescent molecules such as NPs are studied by monitoring the fluorescence intensity fluctuations caused by their diffusion through a small observation volume.^68^ An autocorrelation analysis of these fluctuations yields information on the number of dye molecules per particle, particle concentration, and average diffusion coefficients, which enables the calculation of the hydrodynamic radius of the fluorescent species. Thus, FCS can be used to monitor the eventual disassembly of fluorescently labeled micelles upon dilution or interaction with blood proteins.

Furthermore, if two fluorescent species emitting in different spectral ranges are simultaneously present in the studied system, dual-color fluorescence cross-correlation spectroscopy (FCCS) experiments can also be performed. The amplitude of the cross-correlation function is proportional to the fraction of dual-colored species.^67^ Consequently, the dynamic exchange of unimers between two non-cross-linked micellar solutions, in which different fluorescent dyes are covalently attached to the respective micelle-forming polymers, can be monitored by FCCS after mixing the two solutions. This is because an exchange of unimers will lead to the emergence of a dual-colored species,^69^ whereas an absence of the latter provides evidence for the stability of the micelles. Therefore, FCS experiments were performed to investigate changes in hydrodynamic radii and aggregate formation in plasma or lung surfactant, while FCCS was used to investigate the exchange dynamics of micelles in these body fluids. Besides FCCS, micellar integrity and stability can also be studied by fluorescence resonance energy transfer (FRET), as shown by Hu and Tirelli.^70^

For FCS studies, micelle-forming block copolypept(o)ides were labeled with Alexa Fluor 647 and used to form PM with and without pretomanid and drugs A-D. In an aqueous solution, these micelles displayed an average hydrodynamic diameter of 88-126 nm, consistent with the DLS results (**Table 2**). Most importantly, the hydrodynamic diameter did not change in the presence of serum proteins or lung surfactants, which underlines the pronounced stability of π-π-PeptoMicelles with or without drug loading. For the FCCS analysis, we individually labeled separate batches of pGlu(OBn)_27_-*b*-pSar_182_ at the hydrophilic polymer end group with one of the two fluorescent dyes, Oregon Green 488 and Alexa Fluor 647, and used these different polymers for the preparation of two batches of PeptoMicelles by DC. No unimer exchange between the two systems was detected after incubation in an aqueous solution at room temperature for one week (**Figure S8A**). Even after adding 10 vol% (or 80 vol%) DMSO, a potent solvent for both polymer blocks, no exchange between differently labeled PeptoMicelles occurred after one week at room temperature (**Figure S9**) indicating the high stability of these micelles. By contrast, in previous studies by Schaeffel *et al.*, the addition of tetrahydrofuran (THF) resulted in an increase in exchange dynamics for PM consisting of polystyrene-*block*-poly[oligo-(ethylene glycol) methyl ether methacrylate].^69^

The absence of any detectable exchange of unimers between two separately prepared batches of PeptoMicelles is consistent with high stability and suggests their potential as a long-circulating drug delivery system for hydrophobic drugs. Therefore, we asked whether the micellar stability was altered in the presence of the hydrophobic drug molecules. For this, we prepared micellar drug formulations with drug D, as described above, using the two different fluorescently labeled polymers. The behavior of drug delivery systems in the presence of biological fluids is of particular interest here because stability *in vivo* is a crucial parameter for any therapeutic application. With this in mind, we performed FCCS measurements not only in aqueous solution but also in human blood plasma^71,72^ and in the presence of pulmonary surfactant (Infasurf; **Figure 3**).

**Figure 3.**
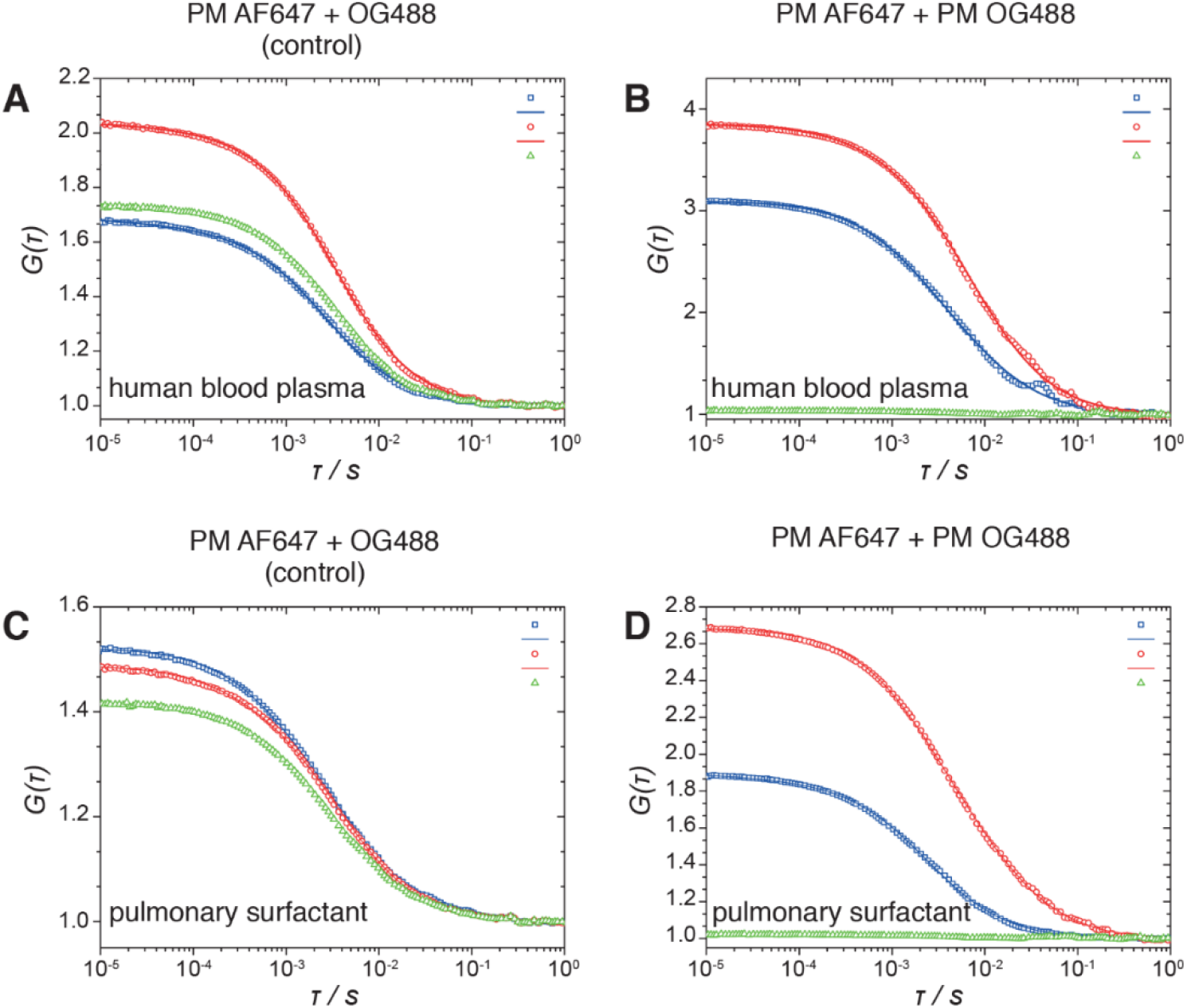
Auto- and cross-correlation functions derived from FCCS measurements of drug D micellar formulations in human blood plasma (**A** and **B**) and pulmonary surfactant (Infasurf; **C** and **D**). Blue: Oregon Green 488 (OG488) experiment (squares) and fit function (line). Red: Alexa Fluor 647 (AF647) experiment (squares) and fit function (line). Green: Cross-correlation of both fluorescence signals. **A** and **C:** Positive control measurements for a drug D NP formulation labeled with both dyes after incubation in the respective medium at 4 °C for one week. **B** and **D:** Measurements were performed after mixing formulations labeled individually with OG488 or AF647 and incubation in the respective medium at 4 °C for one week.

For these FCCS experiments, micellar formulations were diluted with the respective medium and incubated for one week at 4 °C to avoid plasma degradation (expected at 37 °C); all subsequent measurements, however, were performed at 37 °C. Studies of exchange dynamics at higher temperatures were not possible in plasma due to its thermal instability, which induces aggregation of serum proteins. As a positive control for the occurrence of a cross-correlation between the two fluorescence signals, we also analyzed an equivalent micellar drug D formulation made from a diblock copolymer labeled with both dyes. Here, a clear cross-correlation signal could be detected, as shown in **Figures 3A** and **3C**. Like the previous measurements of PeptoMicelles without drug-loading, no dynamic exchange was observed for micellar formulations containing drug D after mixing and incubation in aqueous solution at room temperature one week (**Figure S8B**) or at 60 °C overnight. Strikingly, no occurrence of a cross-correlation function was detected for either the sample incubated in human blood plasma or that incubated in pulmonary surfactant (**Figures 3B** and **3D**). At the same time, the continued presence of the two types of fluorescently labeled PM, which did not change significantly in hydrodynamic diameter after incubation in blood plasma or pulmonary surfactant, was evident from the respective autocorrelation curves. Thus, we concluded that no detectable exchange or aggregation occurs between the individual micelles, with or without drug D, in the media used in this study. Hence, both FCS and FCCS analysis revealed the excellent stability of PeptoMicelles and drug-loaded micellar formulations prepared from the amphiphilic copolypept(o)ide pGlu(OBn)_27_-*b*-pSar_182_ in relevant physiological media (*i.e.*, human blood plasma and pulmonary surfactant) under the conditions evaluated here. Furthermore, storage of the drug-loaded micelles at either room temperature or 4 °C for one year did not change their size distribution or average diameter.

To confirm that this high micelle stability was due to the aromatic groups in the polypeptide side chain in the micelle core, we also synthesized pGlu(O-*tert*Butyl)_32_-*block*-pSar_202_ (not containing aromatic groups and of comparable block length; see **Figures S4** and **S5**), individually labeled separate batches with one of the two fluorescent dyes, and subjected the resulting micellar formulations (not including any drug) to FCS and FCCS analysis under the same conditions as before. In this case, already in the FCS studies, we observed the formation of aggregates in human plasma, indicating reduced stability, presumably due to more interactions of unimers and micelles with serum proteins. These experiments were further supported by FCCS analysis showing reduced stability for micelles based on pGlu(O-*tert*Butyl)_32_-*block*-pSar_202_ compared to the ones based on pGlu(OBn)_27_-*b*-pSar_182_. Apart from the formation of aggregates, we also observed a cross-correlation signal, which strongly indicates the occurrence of exchange reactions between unimers (**Figure S10**). Overall, these findings support our argument that π-π interactions substantially enhance the stability of PeptoMicelles, either in the presence or absence of the pretomanid derivative D.

### *In vivo* evaluation in zebrafish larvae

#### Toxicity/tolerance of the micellar drug formulations

While none of the second generation pretomanid derivatives displayed any cytotoxicity toward mammalian VERO cells, it was crucial to determine if their micellar formulations were well tolerated at therapeutically relevant doses in the zebrafish larvae (although these fish are referred to as an embryo up to day three and a larva thereafter,^73^ we used the term “larvae” throughout this paper). We have reported similar toxicity data for thioridazine and its NP formulation.^74^ The hydrophobic character of the compounds made it impossible to dissolve them without the aid of toxic or poorly tolerated solvents; therefore, only the drug-loaded π-π-PeptoMicelles were used for this initial screen. The toxicity/tolerability was evaluated in healthy uninfected larvae, based on the overall survival of fish (**Figure S12**); in addition, obvious morphological signs of toxicity, *e.g.*, edemas, were recorded.

For the micellar formulations of compounds B, C, and D (at an equivalent drug dosage of 75 ng) and the non-loaded PeptoMicelles (polymer concentration corresponding to drug-loaded particles), the overall survival after eight days was >90%, arguing that none of these compounds nor the PM formulations induced a detectable toxic response. However, the PM formulation of drug A caused high mortality that resulted in all fish being dead by day six post-injection. In addition, most larvae developed large edemas before death, a strong indicator of toxicity. That was surprising because compound A was reported to be non-toxic and well-tolerated in mice at higher doses than tested here (when administered orally),^8^ and the compound was non-toxic toward VERO cells. Therefore, drug A was reexamined at lower doses in micellar formulation or free form, solubilized in a 4:1 mixture of PEG400 and DMSO (tolerated by the fish at low volumes; **Figures S13** and **S14**). At 45 ng in NP formulation, most of the larvae survived for the entire duration of the eight-day experiment (75%). However, even at this dosage, it was still possible to see edemas that were not observed in uninjected zebrafish, and survival was worse for the solubilized free drug (whereas the solvent control was non-toxic at the volumes employed). Hence, we concluded that compound A was not well tolerated in zebrafish and discontinued further assessment.

#### Efficacy of the micellar drug formulations - Blood infection model

Our next objective was to determine whether the remaining PM-encapsulated pretomanid derivatives offered any therapeutic effects *in vivo*. Zebrafish larvae were intravenously challenged with fluorescently labeled (dsRed) *Mm* at day two post-fertilization and then given a single NP dose 24 h post-infection by the same route. **Figure 4B** provides an overview of the timeline associated with treatment studies using the blood infection model. The PM-encapsulated drugs B, C, and D were evaluated head-to-head at an equivalent drug dose of 60 ng, slightly lower than what was used for the toxicity screen. The survival curves and bacterial burdens five days after infection are shown in **Figures 4C and 4D**.

**Figure 4.**
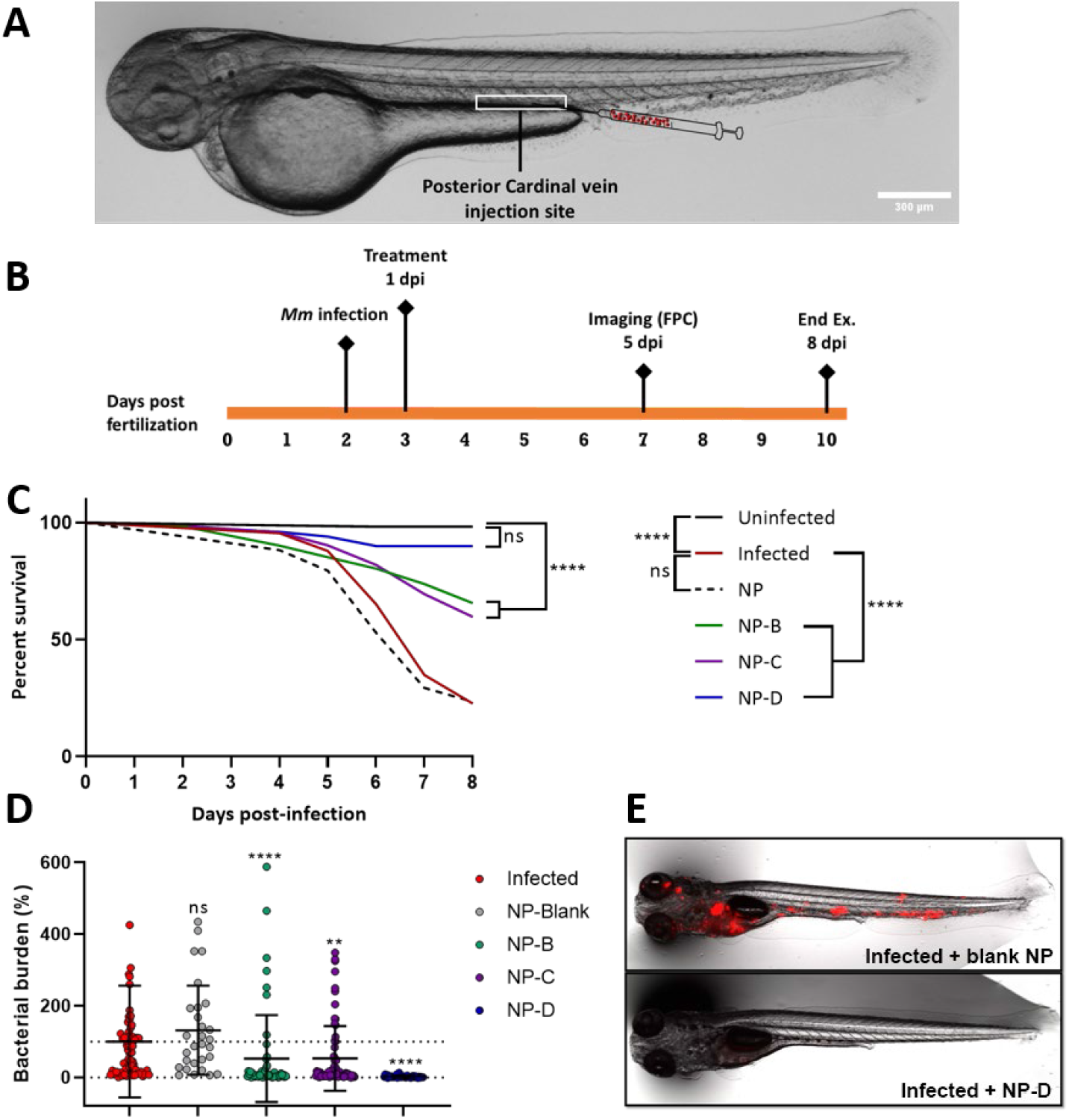
Head-to-head comparison of the therapeutic efficacy of three pretomanid derivative micellar formulations against *Mm* in blood-infected zebrafish larvae. The dosage of each formulation was equivalent to 60 ng of drugs B, C, or D; blank NPs (non-loaded micelles) were used as a control. **A:** WT zebrafish larvae three days post-fertilization. The white box outlines the general injection area for blood infection and treatment. Scale bar 300 µm. **B:** Schematic timeline of the blood infection model. **C:** Survival analysis, N (zebrafish per group at the day of infection) ≥ 34. **D:** FPC analysis of bacterial burden for individual larvae at day five post-infection. Reference group: Infected (untreated control), normalized to 100%. N (zebrafish per group at the day of imaging) ≥ 29. **E:** Representative images from the FPC analysis. On top, an infected zebrafish in the untreated control group, *Mm* in red. Below, an infected zebrafish that had received 60 ng of drug D as a micellar formulation, depicting almost complete elimination of the infection.

As is evident in **Figure 4** for both survival and bacterial burden, compound D was the most efficacious drug in this zebrafish-TB model, consistent with the *in vitro* potency data (**Table 1**). The non-loaded NPs displayed no therapeutic effect. Intriguingly, in previous studies with the TB Alliance, drug D (30 mg/kg) had also exhibited efficacy superior to B and C against *Mtb* in a chronic infection BALB/c mouse model (an eight-week study).^7^ However, for various reasons (including better solubility and lower plasma protein binding), drug C (TBA-354) was selected as the preferred development candidate that was subsequently evaluated in phase I clinical trials.^75–77^

Having confirmed the superiority of drug D in this blood infection zebrafish-TB model, we next wanted to establish a safe and effective therapeutic range for its micellar formulation. For this purpose, two lower doses (37.5 ng and 20 ng) that could be compared to the free drug and two higher amounts (150 ng and 300 ng) were assessed. The results are shown in **Figure 5**.

**Figure 5.**
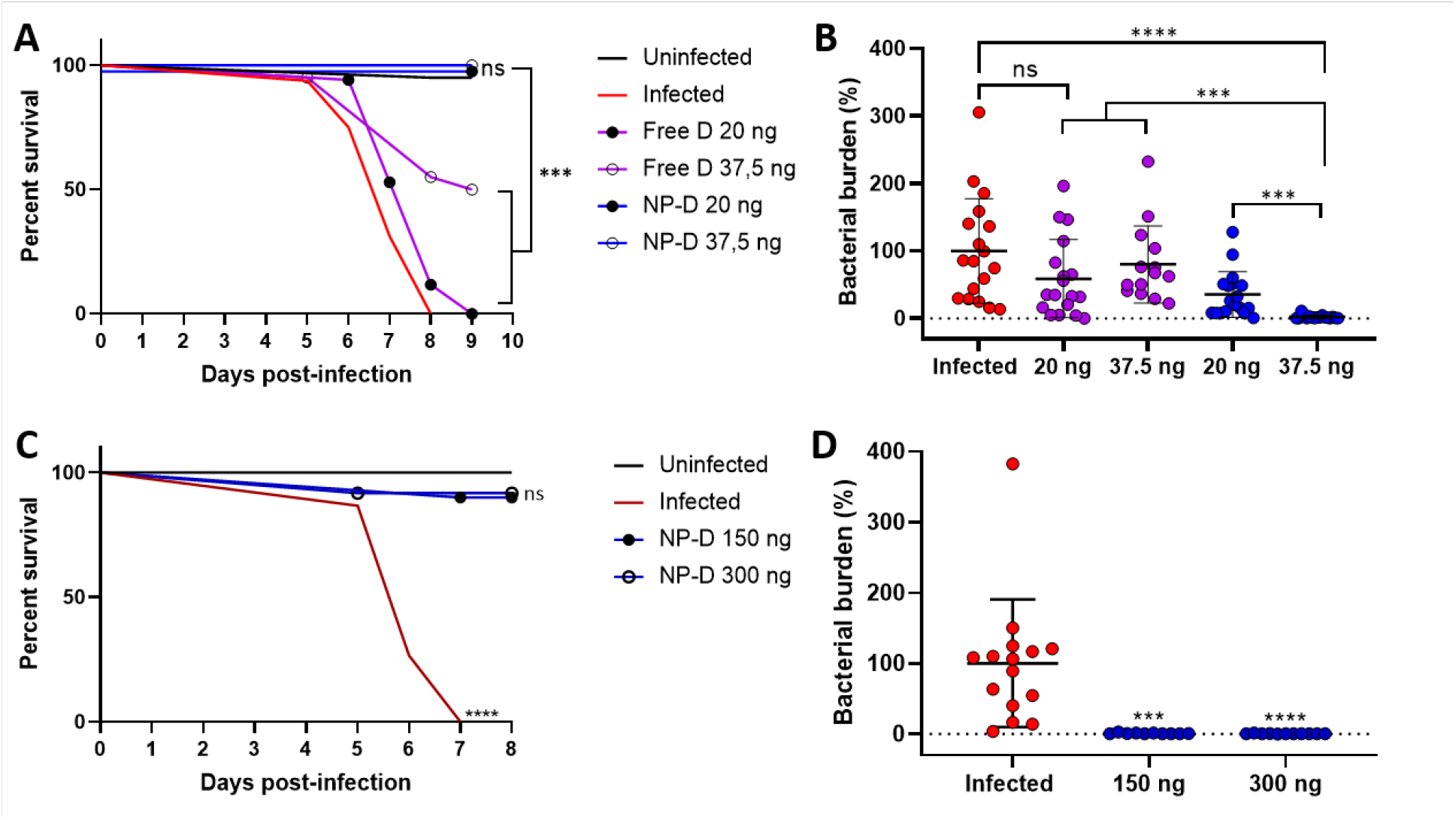
Determination of the therapeutic range of the drug D micellar formulation against *Mm* in blood-infected zebrafish larvae. **A:** Survival analysis at the two low doses tested, 20 ng and 37.5 ng, compared to the free drug. Reference curve: Uninfected (untreated control), n (zebrafish per group at the day of infection) ≥ 16. **B:** FPC analysis of bacterial burden at day five post-infection for the two low doses tested compared to the free drug. Reference column: Infected (untreated control), n (zebrafish per group at the day of imaging) ≥ 14. **C:** Survival analysis at the two high doses tested, 150 ng and 300 ng, respectively. Reference curve: Uninfected (untreated control), n (zebrafish per group at the day of infection) ≥ 10. **D:** FPC analysis of bacterial burden at day five post-infection for the two high doses tested. Reference column: Infected (untreated control), n (zebrafish per group at the day of imaging) ≥ 10.

In this model, the micellar formulation of compound D was safe and effective at doses ranging from 20 to 300 ng (single dose per larva). The highest amount (300 ng) represented an upper boundary, based on a maximum single injection volume of 20 nL and physical limitations regarding the concentration of the formulation itself. The free drug, dissolved in PEG400:DMSO (4:1 ratio), could only be safely administered at the two lowest doses due to vehicle-related toxicity at volumes above 2 nL (**Figure S13**). NP containing drug D at doses of 20 ng and 37.5 ng proved superior to free drug administration in survival analysis and bacterial burden reduction (**Figures 5A and 5B**). Higher NP-D doses (150 and 300 ng; **Figures 5C and 5D**) produced results identical to those for the 37.5 ng dose. Therefore, the latter was considered the minimum effective dose.

#### Circulation time analysis of the lead micellar drug formulation

Due to the excellent therapeutic results, especially with drug D, and the high *in vitro* stability of the micellar NP formulation, we then investigated whether this stability could be maintained *in vivo*, such as in zebrafish larvae. For this purpose, PeptoMicelles with covalently attached Alexa Fluor 647 were injected at 48 h post-fertilization into the posterior cardinal vein (PCV) of zebrafish larvae. Both blank (no drug) PeptoMicelles and drug D-loaded micellar formulations were tested to explore if drug loading interfered with the circulation dynamics. The following time points post-injection were selected for live fluorescent imaging; 5 min (regarded as the 100% level for all NP being in circulation), then 1, 4, 8, 24, 48, and 72 h; in addition, non-injected fish were imaged to represent a zero percent control group. The analyses were performed in ImageJ by the manual option in the *Circulation Time Analysis* macro that we recently published.^78^ The macro analyses were verified by a separate independent evaluation, which resulted in close to parallel circulation dynamics (data not shown). Findings for the drug D formulation and non-loaded micelles are depicted in **Figure 6**.

**Figure 6.**
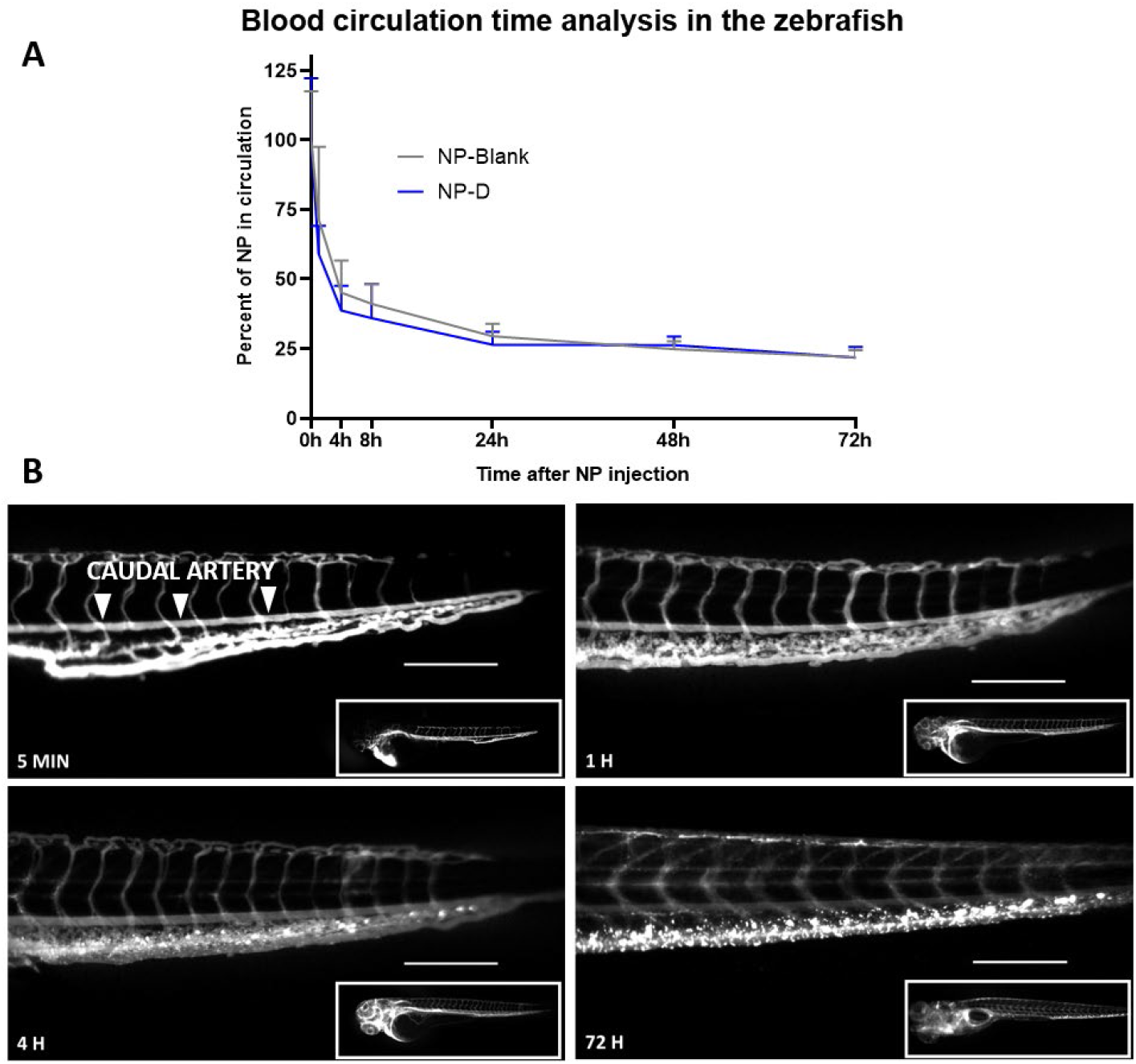
Circulation time analysis for micellar formulations (with or without drug D) in zebrafish larvae. **A:** The particles, with and without drug D, displayed similar circulation percentage versus time profiles, with an initial decline during the first 24 h and then a plateau effect. N (zebrafish per time point) ≥ 5. **B:** Representative images of zebrafish larvae injected with drug D-loaded fluorescent NPs at different time points (5 min, 1 h, 4 h, and 72 h). The decrease in fluorescence of the caudal artery can be seen at time points from 1 h onwards, relative to the 5 min image. The values for fluorescence in the artery were normalized relative to the overall NP fluorescence in the whole fish (inset). Scale bar 200 µm.

Non-loaded PeptoMicelles and drug D-loaded micellar formulations presented comparable results, showing that drug loading did not alter the circulation characteristics of the particles. The circulation-time profile was similar to that of other long-circulating NPs previously tested by us in zebrafish and mice, *e.g.*, PEGylated liposomes and covalently core-crosslinked micelles based on pSar*-b-*pCys(SO_2_Et) copolymers.^78^

#### Granuloma accumulation and efficacy of the lead micellar drug formulation - Neural tube infection model

Injection of *Mm* into the caudal vein of the zebrafish larvae results in small granuloma-like structures that are intimately associated with the vasculature.^51,79^ A significant recent development was injecting the bacteria into the neural tube, an isolated cylinder situated dorsally in the larvae.^80^ This model develops relatively massive granulomas, in which the majority of infected macrophages are undergoing necrosis and cavitation – processes that resemble the situation in human TB. Moreover, this type of granuloma shows extensive angiogenesis and is well vascularized.^47,80,81^ The isolated neural tube also facilitated the potential for two important mechanisms for accumulating NP: the passive EPR effect^34^ and active macrophage uptake, followed by migration to the infection site. Both mechanisms are considered essential strategies for nano-based therapy.^17,47^

As demonstrated above, the π-π-PeptoMicelles were found to be long-circulating. Therefore, we hypothesized that the PeptoMicelle accumulation in granulomas was likely to be augmented by an EPR-like effect. In order to monitor NP interactions and co-localization with granulomas, we first injected wild-type *Mm* (300 CFU) into the neural tube of fli1a:EGFP zebrafish, possessing green fluorescent vasculature, at 72 h post-fertilization. The infection was allowed to progress for three days, and at this time point, the localized neural tube granulomas could be visualized in transmission light by their dark and swollen appearance (**Figures 7B and 7C**). In an earlier study, we found that accumulation of PEGylated liposomes peaked at 4 h, with only a slight increase at 12 h.^47^ Because of this observation, we decided to quantify and compare the accumulation of PeptoMicelles with and without encapsulated drug D at 5 h after NP injection (**Figure 7A**).

**Figure 7.**
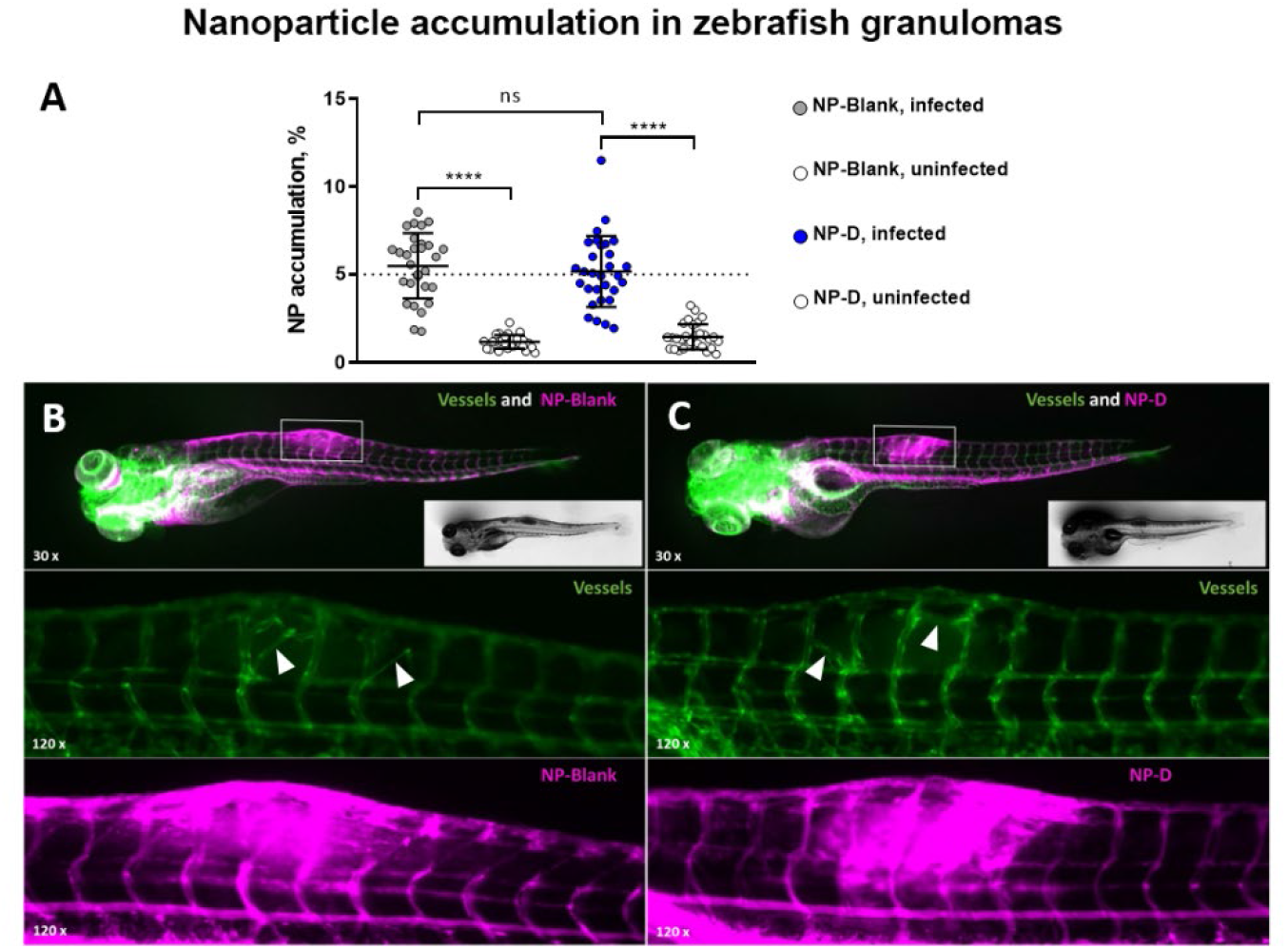
NP accumulation in granuloma-like structures 5 h after injection. **A:** Comparison of PeptoMicelles with and without encapsulated drug D. The background signal represents NP accumulation in an adjacent uninfected area of equivalent size. **B:** Overview of a fish 5 h after injection with PeptoMicelle formulation containing no drug. On top, a 30x image showing both the fluorescent vasculature and NPs, with *Mm* granuloma protruding upward on the dorsal side of the larva (white rectangle). The inset shows a bright field image of the same larva. In the middle, a 120x image showing the fluorescent vasculature (green, EGFP) of the infected area in the neural tube. White triangles point out neovascularization. Below, the same image as above, showing the fluorescent NPs (far-red, AF647). **C:** Equivalent to **B**, but the larva was injected with the drug D-loaded micellar formulation.

About 5% of the injected NP (with and without drug) were found in the granuloma area, a finding consistent with the results we reported earlier for PEGylated liposomes.^47^ The latter study also showed a correlation between zebrafish and mice regarding the accumulation of non-loaded PeptoMicelles of an earlier generation, with lung granulomas of *Mtb*-infected mice exhibiting significant NP accumulation relative to adjacent healthy tissue.

We further decided to utilize the neural tube infection model to directly compare the activity of the micellar NP formulation with that of the free drug (in PEG400:DMSO). Based on the *in vivo* efficacy results above for encapsulated drug D in the blood infection model, a minimally effective dose of 37.5 ng was selected. **Figure 8** depicts the survival curve analysis for this experiment and describes the timeline for the infection model in general.

**Figure 8.**
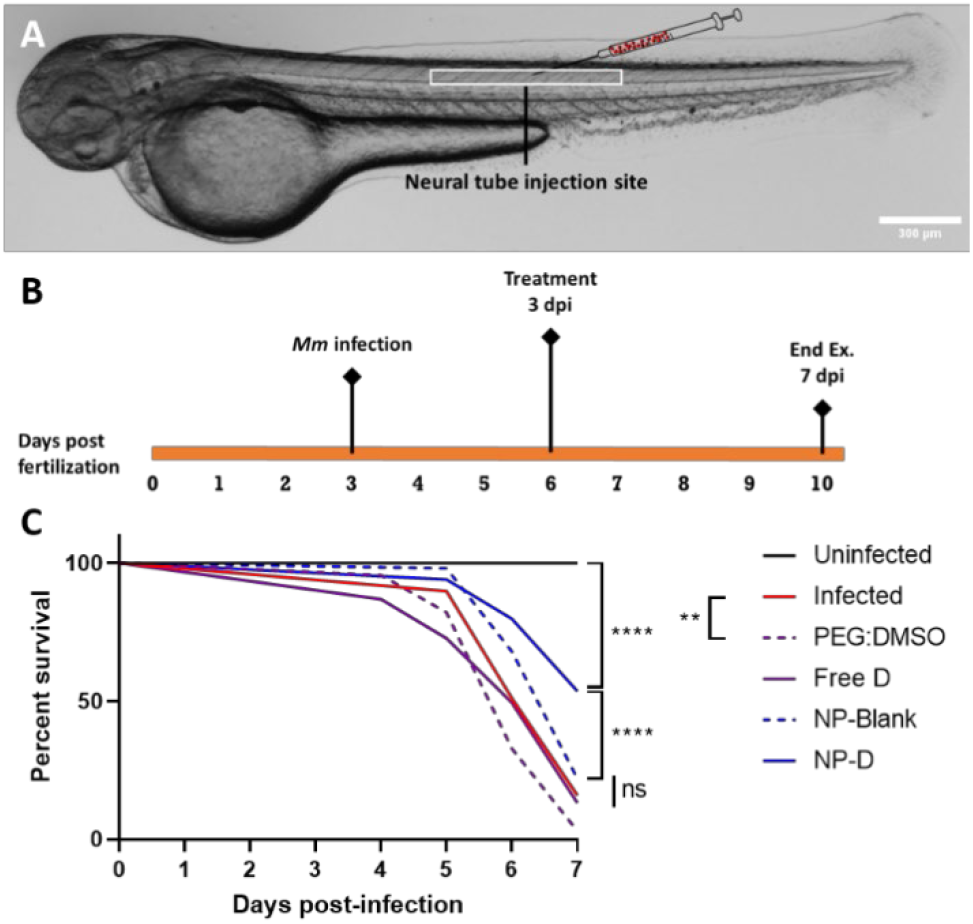
Head-to-head comparison using survival analysis of the therapeutic efficacy of drug D in micellar NP formulation *vs.* drug D solubilized in PEG400:DMSO (4:1 ratio), controls included (PEG400:DMSO and NP-Blank). Zebrafish larvae were infected in the neural tube, and treatment was administered by the posterior cardinal vein. **A:** WT zebrafish larvae three days post-fertilization. The white box outlines the general injection area for neural tube infection and treatment. Scale bar 300 micrometers. **B:** Schematic timeline of the neural tube infection model. **C:** Survival analysis, N (zebrafish per group at the day of infection) ≥ 30.

Fish treated with the free form of drug D (solubilized in 4:1 PEG400:DMSO), vehicle alone (4:1 PEG400:DMSO), or the π-π PeptoMicelles without drug had an almost identical survival curve to the infected control group. In contrast, the corresponding dose of drug D in NP formulation resulted in >50% of fish surviving after seven days post-infection.

### *In vivo* evaluation of the lead micellar drug formulation in mice

Our next objective was to explore the efficacy of this lead micellar formulation against experimental TB in mice. For this purpose, we selected the susceptible C3HeB/FeJ (Kramnik) mouse model because, unlike the commonly used C57BL/6 and BALB/c mice, this mouse strain develops diverse types of lung lesions, including hypoxic and well-structured granulomas in the lungs (with caseous necrotic centers and neutrophilic infiltrates), following low-dose aerosol infection.^82,83^ Hence, it is anticipated that efficacy data in this model may be somewhat more predictive of drug activity in humans.^83^

Two successive mouse experiments were performed to assess the new micellar anti-TB drug formulation. In the first, the micellar formulation of drug D was evaluated alongside non-loaded PeptoMicelles, free drug D dissolved in 4:1 PEG400:DMSO,^84^ and the vehicle alone. (Before commencing this experiment, the safety of multiple doses of 4:1 PEG400:DMSO was confirmed by applying the dosing schedule below to three uninfected mice; no toxic responses or deaths were observed). Drug doses corresponded to 40 mg/kg for a 25 g mouse (the average mouse weight throughout this experiment was 26-27 g), comparable to the minimum efficacious dose in zebrafish but slightly higher than that used in the original eight-week chronic *Mtb* infection study in BALB/c mice.^7^ Dosing commenced 30 days after low-dose aerosol infection with the H37Rv strain of *Mtb*, and all treatments were administered intravenously (via the tail vein) every second day for two weeks. Mice were sacrificed on day 44 post-infection, and organs and blood were removed for plating of tissue samples, histopathological analysis, and cytokine and chemokine measurements.

Our rationale for the above dosing schedule stemmed from the lengthy 24 h half-life in both plasma and lung tissue reported for drug D in CD-1 mice, following oral dosing at 40 mg/kg,^7^ coupled with our expectation that NP formulation would lead to sustained drug release over an extended period. This strategy was subsequently validated by a pharmacokinetic study of both intravenous drug D formulations in uninfected C3HeB/FeJ mice. These new formulations (40 mg/kg) displayed half-lives of 48-55 h and prolonged high drug exposure levels over 48 h, up to 10-fold higher than achieved by oral dosing^7^ and 2-3 orders of magnitude above the MIC value (**Table S1** and **Figure S15**). Nevertheless, processing and analysis of blood samples (see Supporting Information) led to a disintegration of the micelles containing drug D, thwarting our ability to discriminate between micelle-bound versus released drug in plasma.

Results from the CFU analysis of the spleen and lung for the first mouse efficacy experiment are depicted in **Figures 9A and 9B**. Here, free drug D and the micellar drug D formulation (at 40 mg/kg) were equally effective, reducing the bacterial burden by approximately 1.6-1.7 log units (spleen) and 0.8-1.0 log units (lung), compared to untreated control mice. As expected, the non-loaded PeptoMicelles displayed no activity against the infection. The greater efficacy observed in the spleen than the lung in the treatment groups could be explained by differences in the disease pathology between both organs. The spleen is well vascularized, even during pathogenesis, whereas caseous necrotic granulomas in the lung are devoid of blood vessels,^85^ thereby impeding direct access to the infection by nanocarriers or the free drug. In the C3HeB/FeJ mouse, necrotic granulomas tend to occur only in the lung,^82^ although two other lung lesion types can also develop and sometimes predominate.^83^ The poorer therapeutic response to many TB drugs in *Mtb*-infected C3HeB/FeJ mice (compared to BALB/c mice)^82,86–88^ has been definitively linked to the prevalence of large human TB-like necrotic granulomas in some mice, which are notoriously difficult to treat.^87–90^

**Figure 9.**
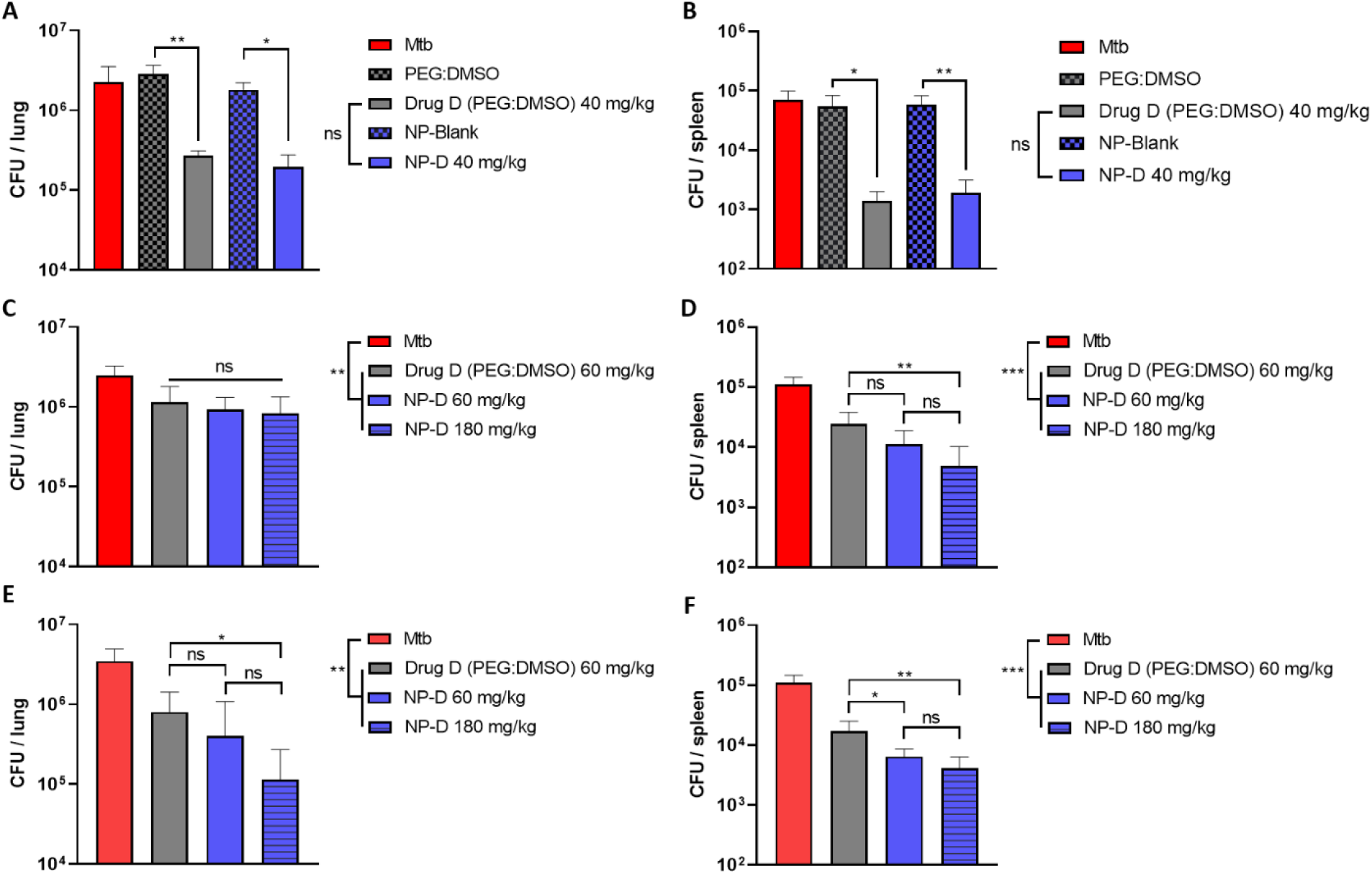
CFU data from lung and spleen samples for two independent efficacy experiments. C3HeB/FeJ mice were aerosol infected with *Mtb* H37Rv, and treatment was administered by i.v. injection on days 30, 32, 34, 36, 38, 40 and 42 post-infection. Mice were sacrificed for plating of tissue samples on d44 post-infection. The CFU data represent the mean and standard deviation from five (A, B) or seven (C-F) mice. **A** and **B:** Experiment 1. Free drug D in 4:1 PEG400:DMSO (40 mg/kg) versus the equivalent drug dose in PeptoMicelles. Both formulations are depicted alongside their corresponding vehicle control (no drug). **C-F:** Experiment 2. Maximum injectable dose of free drug D in 4:1 PEG400:DMSO (60 mg/kg) versus the equivalent drug dose of the micellar formulation and a 3x higher drug dose of the micellar formulation (180 mg/kg). For this experiment, **E** and **F** represent the CFU data from a second dilution of the organ lysates.

Overall, we observed a therapeutic effect from both formulations of drug D, and we postulated that the micellar one might offer a tolerability benefit. Therefore, a second experiment was designed to examine the effects of increasing the dose. We aimed to demonstrate that the nano-formulation was non-toxic and could be administered intravenously at much higher doses than the free drug (using the same route), which was limited by the solubility of the compound (20 mg/mL in 4:1 PEG400:DMSO for drug D) and the known safety threshold for this vehicle.^84^

The second mouse experiment was designed to test the boundaries of the dose level for both drug D formulations while otherwise employing the same dosing schedule and route of administration. We used a slightly higher dose of free drug D (60 mg/kg in 4:1 PEG400:DMSO), an equivalent amount of drug D in the micellar formulation, and a 3-fold higher dose of drug D in the micellar formulation (180 mg/kg). Both 60 mg/kg and 180 mg/kg represented the maximum possible dose levels for free drug D (in 4:1 PEG400:DMSO) and the micellar formulation, respectively. In the latter case, we were constrained by a top injection volume for the mice of 200 µL, a maximal encapsulation percentage of drug in the PeptoMicelles (30%), and a maximum concentration of micellar drug formulation in water (75 mg/mL) to avoid the formation of particulate matter. In the case of free drug, the vehicle volume (75 µL of 4:1 PEG400:DMSO) was the primary limiting factor. Due to viscosity issues, larger injection volumes were too difficult to administer. Higher volumes of this vehicle were also reported to induce toxic responses in mice and hence could not be considered safe for administering multiple drug doses.^84^ (We reconfirmed the safety of 75 µL of 4:1 PEG400:DMSO in three uninfected mice using the same dosing plan, and no toxic responses or deaths were observed.) No adverse effects were noted in either efficacy experiment, and the percentage weight changes for the mice were well within normal thresholds.

In this second mouse study, both pulmonary and splenic CFU data revealed modest but still statistically significant differences between non-treated and treated mice. The 3-fold higher NP-drug dose led to a lower CFU count in the spleen compared to the free drug D treated group (**Figures 9D and 9F**). However, in the case of the pulmonary CFU results, only for one lysate dilution (**Figure 9E**) did the higher NP-drug dose provide significantly greater efficacy than the free drug. Moreover, there was only a weak trend towards better effectivity of the drug D micellar formulation given at 180 mg/kg, compared to the same formulation at 60 mg/kg (CFU differences of 1.6- to 2.3-fold in spleen and 1.1- to 3.4-fold in the lung). Therefore, we also considered histology data obtained from the same experiment (**Figure 10**).

**Figure 10.**
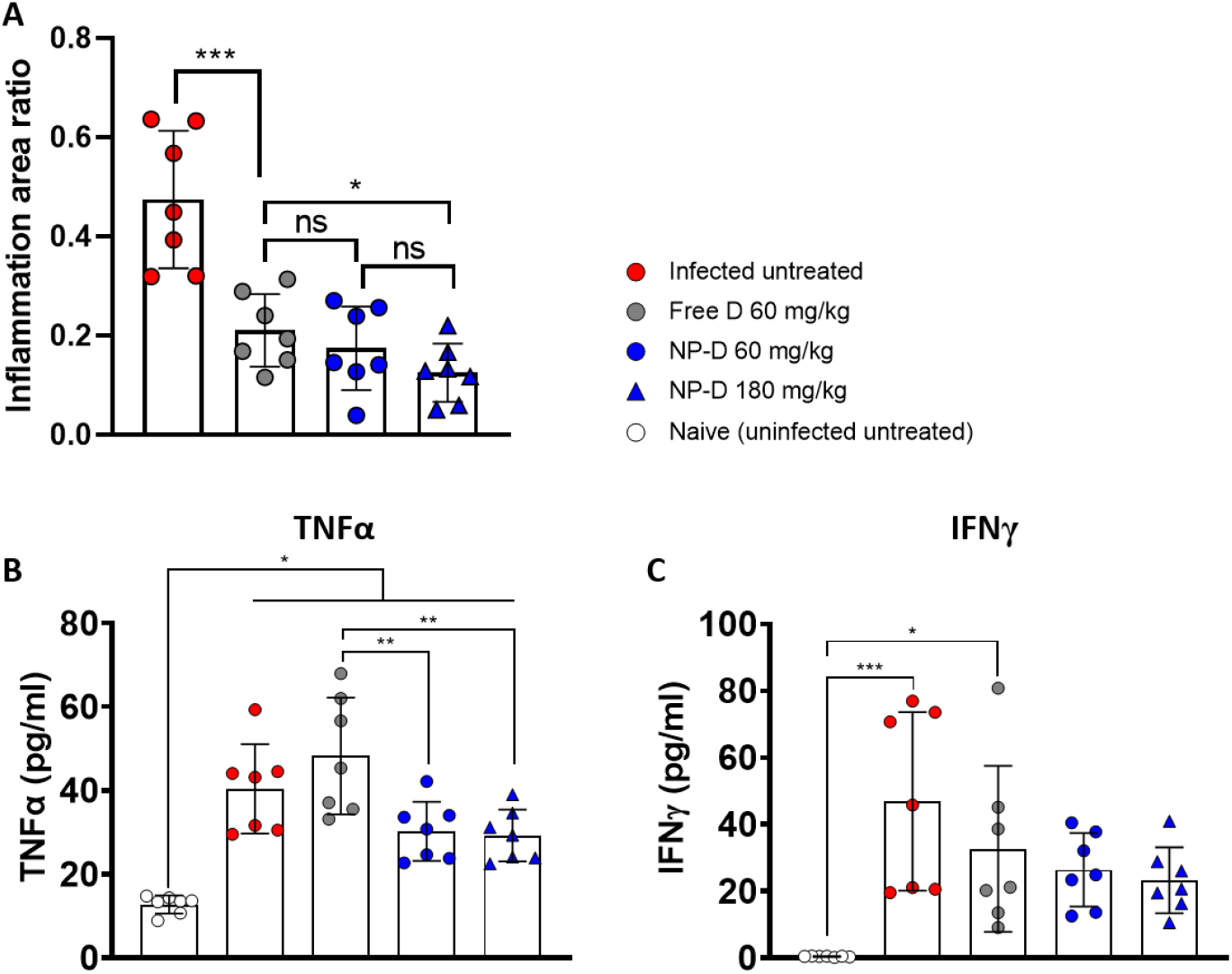
Inflammation data obtained from lung samples for the second mouse treatment study. **A:** Relative ratio of inflamed to non-inflamed area; data derived from lung histological analyses (see the Supporting Information). These data represent the mean and standard deviation from five independent measurements of different lung areas derived from the upper left lung lobe of each mouse (n = 7 mice per treatment group). **B** and **C:** Concentrations of TNFα (**B**) and IFNγ (**C**) in the serum of mice on day 44 after *Mtb* infection. These data represent the mean and standard deviation from pooling two independent replicates per mouse (n = 7 mice per treatment group). Statistical analyses were made using ordinary one-way ANOVA with Tukey’s multiple comparisons test (the mean of each group is compared with that of every other group, and only significant differences are indicated).

Histological analyses of inflamed lung areas were used to correlate inflammation with infectious foci (**Figures S16, S17**). Subsequent quantification revealed that mice treated with either the micellar drug D formulation or free drug D in 4:1 PEG400:DMSO showed markedly reduced areas of inflammatory infiltrates relative to untreated control mice (**Figure 10A**). Furthermore, a maximum dose of the NP drug formulation (180 mg/kg) reduced the inflamed area significantly more than treatment with the free drug at 60 mg/kg. An independent analysis of inflammation was provided by monitoring inflammatory markers in serum collected from all mice at the end of the second experiment. As seen in **Figures 10B** and **10C**, mice treated with the micellar drug D formulation showed on average lower levels of the inflammatory markers IFNγ and TNFα than mice treated with the free drug or infected untreated mice. Data for additional pro-inflammatory markers (**Figure S18**) revealed a similar broad trend of lower levels in NP-treated mice relative to free drug-treated mice.

Returning to the CFU data in this second mouse experiment, the difficulty in detecting a consistent dose-response effect for the micellar formulation of drug D in this model could be due to several factors. First, a two-week dosing period for treating chronically *Mtb*-infected mice is relatively short, and differences between the various experimental groups typically become more apparent after extended dosing periods of four or eight weeks.^7,82^ Second, pretomanid has demonstrated a “maximum bactericidal effect” in human clinical trials, such that doses above a particular threshold value do not lead to any greater efficacy.^91^ A 40 mg/kg dose of the micellar formulation of drug D already delivers exceptional plasma exposure levels over 48 h (**Figure S15**) and significant anti-TB efficacy in the lungs and spleens of treated mice (**Figures 9A** and **9B**). Hence, the higher dose of encapsulated drug D (180 mg/kg) may exceed that required for a maximum bactericidal effect under the dosing schedule employed, resulting in a smaller than expected efficacy difference. Third, there is much greater variability in lung lesion type between mice in the C3HeB/FeJ model than for other mouse models of TB (*e.g.*, BALB/c), leading to a more heterogeneous response to antibiotics.^83,87,88^ This manifests in larger standard deviations for the mean CFU data, reducing the statistical power to distinguish modest differences between the treatment groups.^88^

## CONCLUSIONS

In this study, we developed new micellar drug formulations of four second-generation hydrophobic pretomanid derivatives based on the amphiphilic polypept(o)ide pGlu(OBn)-*b*-pSar. Due to π-π-interactions between aromatic groups in the hydrophobic polypeptide block and with the electron-deficient aromatic systems in the applied drugs, enhanced stability of these micelles was observed, in contrast to micelles without aromatic groups (pGlu(O-*tert*Butyl)_32_-*block*-pSar_202_). The stability of both free and drug-loaded PeptoMicelles under physiological conditions was investigated by FCCS in blood plasma and lung surfactant. This analysis demonstrated the striking stability of the benzyl group-containing polymeric drug carrier system, even in the presence of blood components and natural surfactants. Therefore, this new class of micelles stabilized by π–π interactions was named π-π-PeptoMicelles and further validated *in vivo*.

The selected pretomanid derivatives were non-toxic in mammalian cell cultures and displayed low or submicromolar potencies against *Mm*, with drug D being the best. Evaluation of toxicity and efficacy for the new drug delivery system in a zebrafish larvae model of TB confirmed the safety and superiority of π-π-PeptoMicelles containing drug D over the other encapsulated analogues and the free drug. Measured *in vivo* circulation times for both blank and drug-loaded π-π-PeptoMicelles of several days in zebrafish larvae revealed their “stealth” abilities that could provide the basis for a successful TB therapy through passive accumulation in granulomas. Efficacy studies in the C3HeB/FeJ mouse model revealed therapeutic effects for the micellar formulation of drug D given intravenously at all doses tested, and a 180 mg/kg dose resulted in better therapy than free drug D treatment (60 mg/kg). Although equivalent doses of both formulations gave comparable efficacy using a two-week treatment regimen, the vehicle needed to administer the free drug intravenously (4:1 PEG400:DMSO) is of limited applicability for human use. Moreover, while oral dosing was previously employed for *in vivo* studies of this compound, poor aqueous solubility would still present a challenging hurdle for its further advancement in the absence of a better formulation.

In summary, we present a new micellar anti-TB drug formulation with high stability *in vitro* and *in vivo*, low toxicity and excellent therapeutic efficacy for TB treatment in zebrafish larvae, and promising first results in a more human-like susceptible mouse model for experimental TB. In addition, we have further validated the zebrafish embryo as a powerful intermediate model for drug discovery, where several drugs can be rapidly screened to determine the most promising candidates for assessment in more complex and costly mammalian animal models.

## MATERIALS AND METHODS

### Materials

All reagents and solvents were purchased from commercial suppliers and used as received unless otherwise noted. Before use, THF and *n*-hexane were dried over sodium, and dry DMF was degassed by three freeze-pump-thaw cycles to remove residual dimethylamine. *N,N*-Diisopropylethylamine (DIPEA), triethylamine (TEA), and neopentylamine were dried over sodium hydroxide and fractionally distilled on molecular sieves. Milli-Q water (Millipore) having a resistance of 18.2 MΩ and TOC <3 ppm was routinely employed. Dialysis was carried out using Spectra/Por membranes (Roth) with 3500 g/mol as the nominal molecular weight cut-off. Infasurf (calfactant) was obtained from ONY Biotech (Amherst, USA), while human blood plasma (pooled from six healthy donors and stabilized with EDTA) was provided by the Transfusionszentrale of the Medical Department of the Johannes Gutenberg University Mainz. Borosilicate needles (GC100T-10) from Harvard Instruments were utilized to inject the zebrafish for all experiments.

### Methods

^1^H NMR and diffusion-ordered ^1^H NMR (DOSY) spectra were recorded on a Bruker Avance III HD 400 spectrometer at 400 MHz and room temperature.^92^ All spectra were referenced to the residual solvent signals. Analysis of the ^1^H NMR spectra was performed using the software MestReNova v12.0.0 (Mestrelab Research S.L.).

Analytical hexafluoroisopropanol (HFIP) gel permeation chromatography (GPC) was carried out at a flow rate of 0.8 mL/min at 40 °C with 3 g/L potassium trifluoroacetate added to the eluent.^92^ The GPC system was equipped with a UV detector (Jasco UV-2075 Plus) set at a wavelength of 230 nm and an RI detector (Jasco RI-2031). Modified silica gel columns (PFG columns, particle size: 7 μm, porosity: 100 Å and 1000 Å) were used. Molecular weights were determined by using a calibration with poly(methyl methacrylate) (PMMA) standards (Polymer Standards Service GmbH Mainz) and toluene as an internal standard. The degree of polymerization (DP) of polysarcosine (pSar) was determined by calibration of apparent *M*_n_ against a series of pSar standards characterized by static light scattering to obtain absolute molecular weights.^6158^ Prior to measurement, the samples were filtered through polytetrafluoroethylene (PTFE) syringe filters with a pore size of 0.2 μm. The elution diagram was analyzed with WinGPC software (Polymer Standards Service GmbH Mainz).

Attenuated total reflectance Fourier transform infrared (ATR-FTIR) spectroscopy was performed on an FT/IR-4100 (JASCO Corporation) with an ATR sampling accessory (MIRacleTM, Pike Technologies).^92^ The IR spectra were analyzed with the software Spectra Manager version 2.02.05 (JASCO Corporation); 16 scans were performed per measurement.

Single-angle dynamic light scattering (DLS) and zeta potential measurements were made on a Zetasizer Nano ZS (Malvern Instruments Ltd., Worcestershire, UK) at an angle of 173° and a wavelength of 633 nm at 25 °C.^92^ Three measurements were performed per sample at concentrations between 0.1-0.3 mg/mL in 10 mM NaCl solution, and size distribution (intensity-weighted) histograms were calculated based on the autocorrelation function of samples, with automated attenuator adjustment and multiple scans (typically 10-15 scans). Prior to recording these data, samples were filtered through 0.2 μm GHP membrane filters (Pall Corporation, Port Washington, USA). Disposable polystyrene cuvettes (VWR, Darmstadt, Germany) were used for size determinations and disposable folded capillary cells (Malvern Instruments Ltd., Worcestershire, UK) for zeta potential measurements. The data were analyzed with Malvern Zetasizer Software version 7.12.

Melting points were measured using a Mettler FP62 melting point apparatus at a heating rate of 1 °C/min.

#### Transmission electron microscopy (TEM)

The staining and embedding were performed using freshly filtered solutions. Micellar NP formulations, diluted to 100 µg/mL, were adsorbed onto carbon-coated (300 mesh) copper grids for two minutes floating, followed by 2×1 min washing steps with H_2_O. Samples were quickly passed on one drop of 2% uranyl acetate (UA) directly, followed by 30 seconds on a 1% UA drop and 2×1 min washing, as before. For embedding, the grids were then transferred to a drop of 0.4% UA/1.8% methylcellulose on ice, followed by 2 min incubation on another drop of 0.4% UA/1.8% methylcellulose, followed by pickup in loops and drying. Microscopy was performed using a JEOL JEM 1400 TEM equipped with a TWIPS camera.

#### Cryo TEM

About 5 µL of undiluted drug D-loaded micellar NP formulation was applied to a holey carbon film on a 300 mesh grid, blotted and plunge-frozen in liquid ethane. Microscopy was carried out on a JEOL JEM 1400 equipped with a TWIPS camera, using a GATAN cryo-holder.

Fluorescence cross-correlation spectroscopy (FCCS) experiments were performed using a commercial setup LSM 880 (Carl Zeiss, Jena, Germany). For excitation of the OG488 and AF647, an argon ion laser (488 nm) and a He/Ne-laser (633 nm) were used simultaneously. A water immersion objective with a high numerical aperture (C-Apochromat 40x/1.2 W, Carl Zeiss, Jena, Germany) was employed to focus the excitation light into the sample and to collect the fluorescence. The fluorescence was then passed through a confocal pinhole and directed into a spectral detection unit (Quasar, Carl Zeiss). In this unit, the emission is spectrally separated by a grating element on a 32 channel array of GaAsP detectors operating in a single photon counting mode. The emission of OG488 was detected in the spectral range 500-553 nm and that of AF647 in the range 642-696 nm. These arrangements formed two overlapping confocal observation volumes, *V*_b_ and *V*_r_, that superimpose to a common observation volume *V*_br_. The observation volumes were calibrated using Alexa Fluor 488® and Atto Fluor 643® as reference dyes with known diffusion coefficients.

After the respective incubation time, the micelles were further diluted (see below) and poured into eight-well polystyrene-chambered coverglass (Laboratory-Tek, Nalge Nunc International, Penfield, NY, USA) mounted in a microscope incubator (PM 2000 RBT, Pecon, Erbach, Germany).

For each solution, a series of twenty measurements with a total duration of 3 min was performed. The experimental auto- and cross-correlation curves were fitted with the theoretical model function for an ensemble of freely diffusing fluorescence species.^67^ The fits yielded the diffusion times of the fluorescent species from which the respective diffusion coefficients were evaluated. Finally, the hydrodynamic radii (*R*_h_) were calculated (assuming spherical particles) by using the Stokes-Einstein relation:

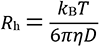

In this equation, *k*_B_ is the Boltzmann constant, *T* is the temperature, *D* is the diffusion coefficient, and *η* is the viscosity of the solvent.

The amplitudes of the autocorrelation curves *G*_b_*(0)* and *G*_r_*(0)* are reversely proportional to the average numbers of “blue” and “red” species in the observation volume, and the amplitude *G*_br_*(0)* of the cross-correlation curve is proportional to the fraction of dual-labeled species with *G*_br_*(0)* = 1, meaning that no dual-labeled species is present.

### Syntheses

Synthesis of sarcosine (Sar) N-carboxyanhydride (NCA), γ-benzyl-L-glutamate (Glu(OBn)) NCA and γ-tert-butyl-L-glutamate (Glu(OtBu)) NCA. NCAs were synthesized as reported.^42,93^

*Synthesis and fluorescent labeling of pGlu(OBn)_27_-*b*-pSar_182_*. The synthesis of pGlu(OBn)-*b*-pSar was adapted from Birke et al.^42^ and modified (see the Supporting Information).

*Synthesis and fluorescent labeling of pGlu(O*t*Bu)_32_-*b*-pSar_202_*. The syntheses are described in the Supporting Information.

*Synthesis of test compounds.* Pretomanid and analogues A-D were synthesized according to previously reported protocols.^6–8,94^ Larger quantities of analogue D for the mouse studies were obtained *via* a modified procedure (see the Supporting Information).

### Nanoparticle preparation

#### Dual centrifugation (DC)

The preparation of π-π-PeptoMicelles was performed similarly to that previously described.^47^ A total of 200 μL (corresponding to 5 mg of block copolymer) of a 25 mg/mL stock solution of pGlu(OBn)_27_-*b*-pSar_182_ in chloroform was added to a 0.2 mL PCR vial, and the chloroform was evaporated overnight. The next day, 50 μL sterile water and 80 mg ceramic beads (SiLibeads ZY-S, 0.3-0.4 mm) were added to the vial. After the polymer was allowed to swell for 4 h, DC was performed for 30 min at 3500 rpm with a dual asymmetric centrifuge (SpeedMixer DAC 150.1 CM, Hauschild & Co.KG). Following centrifugation, the slightly turbid solution was separated from the beads with an Eppendorf Pipette, and the vial was rinsed three times with 50 μL sterile water or PBS at a time. For the preparation of drug-loaded micelles, 3.5 mg polymer and 1.5 mg drug were used in the same DC protocol. For obtaining fluorescently-labeled micelles, 1 mg of the non-functional polymer was replaced by 1 mg of polymer labeled with the respective dye (Oregon Green 488 or Alexa Fluor 647). PM derived from pGlu(O*t*Bu)_32_-*b*-pSar_202_ were prepared under the same conditions as those from pGlu(OBn)_27_-*b*-pSar_182_ to ensure comparability of results.

#### Sample preparation for FCCS measurements

Prior to FCCS analysis, samples were diluted 100-fold (corresponding to a total concentration of 0.25 mg/mL) with the respective medium (water, water + 10% DMSO, human blood plasma, or pulmonary surfactant solution) and incubated for one week at room temperature (for water and 10% DMSO/water) or 4 °C (for human blood plasma or pulmonary surfactant). Directly before the measurement, samples were warmed up to 37°C and diluted 10-fold to obtain dye concentrations (ca. 250 nM) suitable for FCCS measurement. Usually, the respective medium was used for dilution. Only in the case of pulmonary surfactant was dilution with water necessary.

### *In vitro* drug testing

#### Bacterial strains and plasmids

The following bacterial strains and plasmids were used throughout this study: *M. marinum* ATCC BAA-535 wild-type isolate M (with and without pMSP12::dsRed2, resistant to kanamycin, chromosomally integrated) and *M. tuberculosis* H37Rv. *Mm* were grown at 30 °C and *Mtb* at 37 °C in Difco Middlebrook 7H9 broth supplemented with 10% ADC enrichment medium and 0.05% tyloxapol. For dsRed-*Mm*, 40 µg/mL kanamycin was added to the media for antibiotic selection.

#### Resazurin Microtiter Assay (REMA)

The compounds were kept as frozen (−20 °C) stocks dissolved in pure DMSO. Test compounds were thawed at room temperature, added 1:100 in 7H9 media supplemented with 10% OADC to the topmost well of a transparent 96-well microtiter plate (VWR), and serially diluted (2-fold) to the end of the plate. M strain *Mm* and H37Rv *Mtb* were grown without antibiotic selection to mid-log phase, OD_600_ adjusted to 0.8, and diluted 1:100 (*Mtb*) or 1:200 (*Mm*) in PBS. Bacterial inoculum was aliquoted into the wells, ∼10^3^ and ∼10^4^ CFU/per well for *Mm* and *Mtb,* respectively, and plates were incubated at 30 °C (*Mm*) or 37 °C (*Mtb*). Three (*Mm*) or six (*Mtb*) days after plate preparation, 10 μL of 0.03% Resazurin sodium salt solution was added to every well, and plates were incubated for a further 48 h. The MIC against *Mtb* was defined as the lowest drug concentration that prevented a color change from blue to pink. The MIC_90_ against *Mm* was the lowest compound concentration that prevented at least 90% of bacterial growth. These latter values were determined by recording the fluorescence in each well using a microplate reader (BioTek), subtracting background, and calculating the results as a percentage of mean data for untreated controls by Excel. Recorded MIC and MIC_90_ values were the averages of at least three independent determinations.

#### Drug cytotoxicity testing

Compound cytotoxicity was evaluated in VERO (kidney, African green monkey; ATCC CCl-81) cells. VERO cells were cultured in Dulbecco’s modified Eagle’s medium (DMEM) supplemented with 10% fetal bovine serum (FBS) and 2 mM glutamine (Lonza). Cells were dislodged, centrifuged, and resuspended in fresh medium before being dispensed into 96-well plates at 20 000 cells/well (100 µL/well). Test compounds in two-fold serial dilution were added, starting from 200 µM, and cells were incubated for 72 h. Next, a cellular viability test was performed using CellTiter-Glo® Luminescent Cell Viability Assay (Promega), which enumerates how many cells are alive and healthy, based on quantifying the ATP present. From these results, IC_50_ values were calculated (defined as the drug concentration required to give 50% cellular growth inhibition *in vitro*). Cells were also inspected under a light microscope. Selectivity Index data (SI= IC_50_ /bacterial MIC) were calculated using the measured MIC values against *Mm* and *Mtb*.

### Zebrafish experiments

#### Zebrafish Larva Husbandry

Zebrafish larvae were kept in Petri dishes containing larva water^95^ supplemented with phenylthiourea (0.003%, Aldrich) to inhibit melanization. A constant water temperature of 28.5 °C was employed, which is considered optimal for development.^73^ Zebrafish maintenance and experiments were in full compliance with ethical standards and legislation for animal research in Norway. All activity involving zebrafish larvae was approved and overseen by the Norwegian food and safety authority.

#### Zebrafish microinjections

The microinjection protocol that was used was adapted from Cosma and colleagues.^95^ A pipette puller (P-97, Sutter Instruments) was used to prepare the borosilicate needles (GC100T-10, Harvard Instruments) required for the microinjection procedure. The needle was then mounted onto a Narishige MN-153 micromanipulator that enabled fine-tuned movements and connected to an Eppendorf Femtojet Express pump. Before injection, zebrafish larvae were anesthetized by immersion in a tricaine bath (Finquel; 0.02% in larva water). During the injection procedure, the larvae were placed on a Petri dish lid comprising a solidified solution of 2% agarose in Milli-Q water.

#### Infection and treatment of zebrafish

The infections were performed 48 h (PCV) or 72 h (NT) post-fertilization. Approximately 500 CFU (10^8^ CFU/mL, 5 nL) of *Mm* (pMSP12::dsRed2) were injected into the posterior cardinal vein (PCV) or 300 CFU (3 nL) of *Mm* (wild-type) into the neural tube (NT). The larvae were allowed to recover before administering the appropriate treatment formulation *via* the i.v. route. The times of treatment (single dosing) were 24 h (PCV infection) or 72 h (NT infection) after infection. Different doses were administered by adjusting the injected volume or varying the solution concentrations of the treatment formulations. The free form of the drug was dissolved and injected in a 4:1 mixture of PEG400 and DMSO.

#### Zebrafish survival study; infection/treatment and toxicity

One hour after the treatment injection, the zebrafish larvae were controlled, and only live fish were included in the experiment. The larvae were controlled daily, all groups at the same time, and dead and dying larvae were removed.

#### Zebrafish fluorescent pixel count

Zebrafish larvae were imaged at different times, depending on the purpose of the study (bacterial burden or NP accumulation), using a Leica DFC365FX stereomicroscope with a 1.0X planapo lens. For determining bacterial load, zebrafish larvae were imaged on day five post-infection, and images of the whole fish were obtained (30X magnification). For quantifying the fluorescent pixel count (FPC) corresponding to the bacterial burden (*Mm*-DsRed), a customized macro in ImageJ was applied (compatible with *.lif format and *.tif images). The macro first provided for thresholding (adjustable) of the dsRed signal to remove background noise and autofluorescence and define detected Regions of Interest (ROIs). The value used for the analysis was the Raw Integrated Density (RawIntDen), which represented the total sum of the pixel values in the ROIs for each image. The results were stored in *.txt files for each experiment. Data were normalized using GraphPad Prism (version 8), with the average values of the uninfected- and infected-untreated control groups being set to 0% and 100%, respectively.

For quantifying the accumulation of fluorescently labeled NPs in neural tube granulomas, images of the whole fish (30X magnification) were obtained 5 h after injection, four days post-infection. The analysis was performed using the software ImageJ. The fluorescence relative to the area of the granuloma was calculated using two defined ROI (rectangle tool), one large for the whole fish and one small for the granuloma. The same approach was used to calculate the fluorescence relative to the area of uninfected tissue adjacent to the granuloma.

#### Circulation time in zebrafish

AB Wild-type zebrafish larvae were used; the experiment was initiated 48 h post-fertilization. The procedure for imaging and circulation time analysis is described in the paper of Dal et al.^78^ Larvae without visible blood flow were not analyzed.

#### Statistics

Statistical analyses were made using GraphPad Prism (version 8) software. For the FPC analysis, a D’Agostino & Pearson normality test was used to check for Gaussian (normal) distribution within the individual groups. Data showing this pattern were analyzed by one-way Anova (Dunnett’s multiple comparisons test). If no Gaussian distribution was found, data were analyzed by a Kruskal-Wallis test (Dunn’s multiple comparisons test). The survival results were analyzed using the log-rank (Mantel-Cox) test. The significance level (P-value) is indicated as * < 0.05, ** < 0.01, *** < 0.001 and **** < 0.0001.

### Mouse experiments

#### Husbandry

Six to eight-week-old female C3HeB/FeJ mice (Jackson Laboratories, USA) were used for the experiments. Their housing consisted of individually ventilated cages containing filters, located in a specific pathogen-free environment of the Biological Safety Laboratory (BSL) 3. Mice were supplied with adequate food and water from the Research Center Borstel, Germany. All animal experiments were performed according to the German animal protection law and approved by the Ethics Committee for Animal Experiments of the Ministry for Agriculture, Environment and Rural Areas of the State of the Schleswig-Holstein, Germany, under the license V 244-34653.2016 (63-5/16).

#### Bacterial culture

*Mycobacterium tuberculosis* strain H37Rv (*Mtb*) was used for the aerosol infection of mice. *Mtb* was grown in Middlebrook 7H9 broth (BD Biosciences) supplemented with OADC (Oleic acid, Albumin, Dextrose, Catalase) enrichment medium (BD Biosciences). Bacterial cultures were harvested, and aliquots were frozen at -80 °C for later use. Viable bacterial counts were performed by plating the serial dilutions into Middlebrook 7H11 agar (BD Biosciences) plates and incubating them at 37 °C for 21-28 days.

#### Mtb aerosol infection

C3HeB/FeJ mice were infected with the virulent *Mtb* H37Rv strain by the aerosol route at day 0. For aerosol infection of mice, previously prepared frozen *Mtb* stocks were thawed and resuspended, using a 1 mL syringe with a 27G3/4 gauge to achieve a uniform mycobacterial suspension. The bacterial suspension was then diluted in sterile distilled water in a volume of 6 mL to deliver 100 mycobacteria to each animal. A total of 5.5 mL of the diluted bacterial suspension was used for the infection, while the remaining 500 μL was serially diluted until 10^-5^, plated in Mycobacteria 7H11 agar plates, and incubated for 3-4 weeks at 37 °C. Aerosol infection of the mice was performed using a special approach where animals were kept inside a specialized metal cage and placed inside the aerosol chamber (GlasCol) to adapt a natural entry of mycobacteria. A total of 5.5 mL of the bacterial suspension was transferred to the nebulizer connected to the aerosol chamber, and the aerosol process started. Inside the chamber, animals were exposed to low dose *Mtb* aerosol by regulating mainstream airflow (around 1.68 m^3^/h) and the compressed airflow (approximately 0.28 m^3^/h) for nebulizations.

#### Antibiotic treatment

In the first experiment, groups of five mice were administered the respective formulations every second day for two weeks (**Table 3**), seven treatments per mouse in total. In the second experiment, groups of seven mice were employed and were administered the respective formulations using the same dose schedule as above (**Table 4**). At the conclusion of the dosing period in each experiment (day 44), animals were sacrificed, and lungs and spleens were removed for CFU determination and histopathological analysis.

**Table 3.** Overview of the different treatment formulations, doses, and injection volumes used in the first mouse efficacy study.

| **Formulations** | **Dose per injection** | **Volume per injection** |
| --- | --- | --- |
| Infected (terminated at d44) | - | - |
| NP-Blank<br>(25 mg/mL polymer) | 135 mg/kg polymer | 135 μL |
| NP-D<br>(25 mg/mL total, 30% drug) | 40 mg/kg drug D<br>95 mg/kg polymer | 135 μL |
| PEG400:DMSO (4:1 ratio) | 1805 mg/kg PEG400<br>440 mg/kg DMSO | 50 μL |
| Free D (PEG400:DMSO)<br>(20 mg/mL drug)<br>(Solvent 4:1 ratio) | 40 mg/kg drug D<br>1805 mg/kg PEG400<br>440 mg/kg DMSO | 50 μL |

**Table 4.**
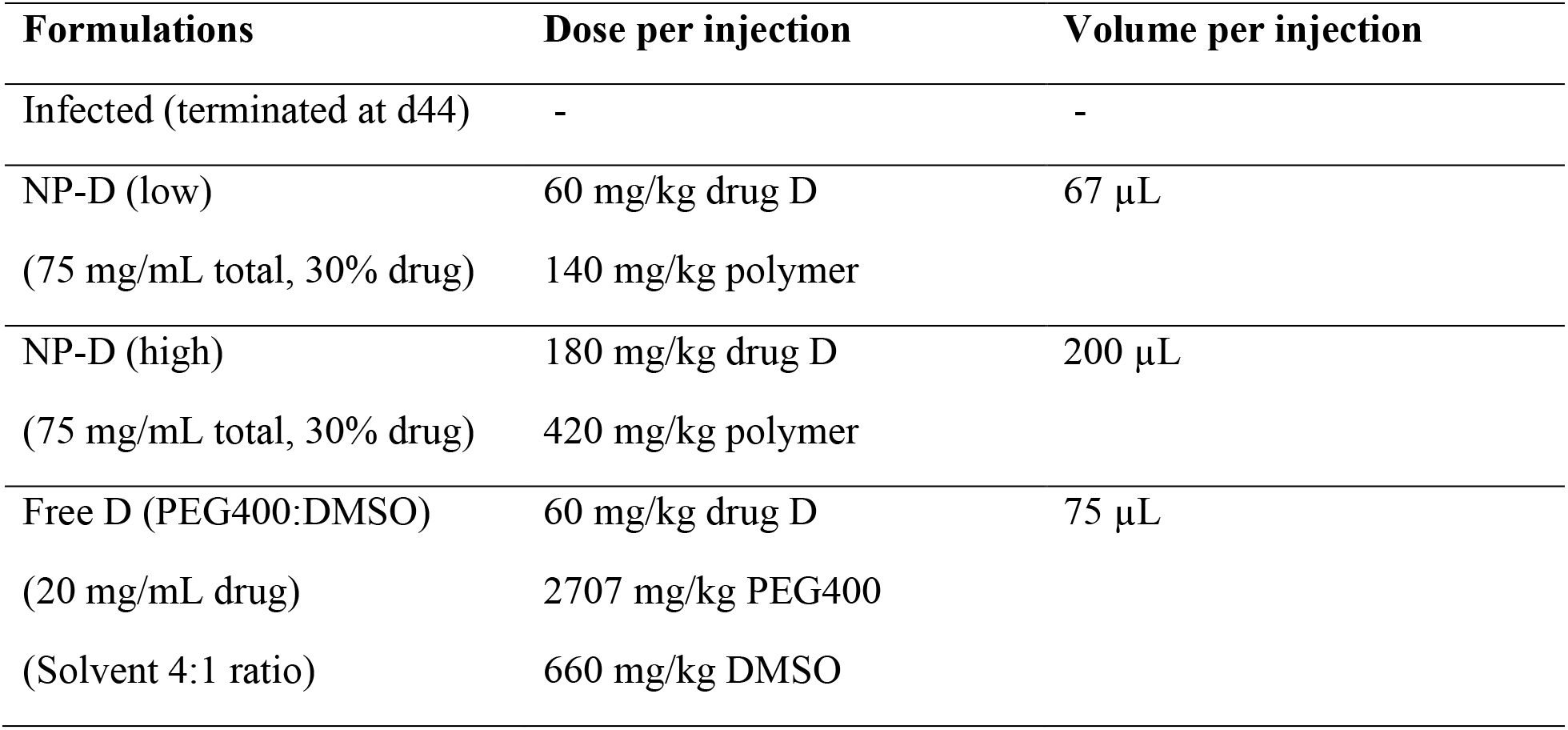
Overview of the different treatment formulations, doses, and injection volumes used in the second mouse efficacy study.

#### Colony-forming unit (CFU) assay

After animals were sacrificed, bacterial burdens in the lungs and spleens were evaluated. Whole organs were harvested, weighed, and mechanically ground in 1 mL WTA (water:Tween-80 (0.01%): albumin (0.05%)) buffer inside the Whirl-pak plastic bag, using a 50 mL falcon tube and Petri dish. Organ homogenates were 10-fold serially diluted in WTA buffer, and 100 μL were plated onto Middlebrook 7H11 agar plates using glass rods and incubated at 37 °C. After 21-28 days, mycobacterial colonies were counted.

#### Quantification of inflammation

Lung tissue was preserved with 4% PFA, cryoprotected using increasing concentrations of sucrose (final concentration 20%) in PBS for 24 h, embedded in Tissue-Tek O.C.T Compound (Sakura Finetek, USA), and flash-frozen in isopentane. Subsequent cryo-sections were stained with H&E and analyzed by bright-field microscopy. Quantification of inflamed areas of the lungs was performed with cell Imaging Software for Life Sciences Microscopy (cellSens Ver.2.2, OlympusBX41). Inflamed and non-inflamed areas were framed manually. Computed data were converted into Excel files and used for quantification. Data were collected from five independent measurements from different lung areas derived from the upper left lung lobe of each mouse (n=7).

#### Multiplex cytokine assay

Blood was collected under terminal anesthesia from the vena cava caudalis (using a 0.4 mm x 19 mm 27 G cannula and a 1 mL syringe) into tubes containing a clot activator (Sarstedt AG &Co. KG., Nümbrecht, Germany) and stored at -4 °C overnight. Mice sera were transferred to 0.22 μm centrifuge filter tubes (Costar., New York, USA), spun at 4 °C for 5 min (8000 rpm), and stored at -20 °C. The sera concentrations of various cytokines and chemokines were determined using a customized U-PLEX Biomarker Group 1 (mouse)-5 spot assay (Meso Scale Discovery, Inc., Rockville, USA). Similar to a sandwich ELISA, the plates were coated with the linker-coupled biotinylated capture antibody solution and incubated overnight at 4 °C. Unbound linker-antibody solution was washed off the plates with 0.05% v/v Tween-20 in DPBS. Serum samples and standards were added in duplicates to designated wells on the plates, incubated for 2 h at room temperature, and washed with 0.05% v/v Tween-20 in DPBS. Detection antibodies conjugated to electrochemiluminescent labels were added to the plates and incubated for 1 h at room temperature. Finally, the plate was mounted into the Meso™ QuickPlex™ SQ 120 instrument, which supplied a voltage to the in-built carbon electrodes on the bottom of the plates, causing the electrochemiluminescent labels bound to the analyte to emit light. The intensity of emitted light was proportional to the amount of each analyte present in the sample measured by the device. The signals of the samples were compared against those of the standard curve. The standard curve was prepared by 4-fold serial dilutions of the calibrator standard 1 till standard 8 to achieve the highest and lowest points in the standard curve, respectively. Data acquisition was performed using the Discovery Workbench® version 4.0 software (Meso Scale Discovery, Inc., Rockville, USA), and data analysis was performed using GraphPad Prism software version 9.0.1 (GraphPad Prism Software Inc., California, USA).

#### Statistics

Statistical analyses were made using GraphPad Prism (version 8) software. Unless otherwise noted in the text, the CFU and inflammation data were evaluated by one-way analysis of variance (ANOVA), and overall differences were considered significant at the 5% level. Differences between groups were further evaluated by Welch’s t-test, and differences were considered significant at the 5% level. The significance level (P-value) is indicated as * < 0.05, ** < 0.01, *** < 0.001 and **** < 0.0001.

## ASSOCIATED CONTENT

### Supporting Information

Additional methods, synthetic and characterization data, drug release profiles, supplementary FCCS and zebrafish data, mouse pharmacokinetic data, and more histology and cytokine data from the second mouse experiment (PDF).

## AUTHOR INFORMATION

### Notes

The authors declare no competing financial interest.

## Supporting information

Suppporting information file

## ACKNOWLEDGMENTS

This work was financially supported by the Norwegian Research Council grants Frimedbio (275873) and Bedrehelse (273319) and the Max Planck Graduate Center with the Johannes Gutenberg-Universität Mainz (MPGC). We thank the Oslo IBV EM Facility (Head N. Roos) and the Imaging Facility (Head O. Bakke) for excellent support. UES and NR thank the Leibniz Association, the European Union (NAREB) and the VDI-VDE (anti-TB) for funding. M.B. acknowledges support by the German Research Foundation (DFG) under the CRC 1066-2/-3.

