## Supplementary material for "Π-Π Interactions Stabilize PeptoMicelle-Based Formulations of Pretomanid Derivatives Leading to Promising Therapy Against Tuberculosis in Zebrafish and Mouse Models": Suppporting information file

### Table of Contents

1. Synthesis, characterization, and fluorescent labeling of pGlu(OBn)$_{27}$-*b*-pSar$_{182}$ and pGlu(O*t*Bu)$_{32}$-*b*-pSar$_{202}$ (pp S3-S8, **Figures S1-S5**)

2. Determination of encapsulation efficiency and drug release profiles (pp S9-S12, **Figures S6-S7**)

3. Additional fluorescence cross-correlation spectroscopy (FCCS) data (pp S13-S14, **Figures S8-S10**)

4. Improved synthetic route for the larger-scale synthesis of pretomanid analogue D (pp S15-S16)

5. Additional zebrafish (ZF) data (pp S17-S19, **Figures S11-S14**)

6. Pharmacokinetic studies in mice (pp S20-S21, **Table S1** and **Figure S15**)

7. Histological analysis of lung sections and analysis of inflammation markers in serum from each experimental group in the second mouse experiment (pp S22-S25, **Figures S16-S18**)

8. References (p S26)

### 1. Synthesis, characterization, and fluorescent labeling of pGlu(OBn)$_{27}$-*b*-pSar$_{182}$ and pGlu(O*t*Bu)$_{32}$-*b*-pSar$_{202}$

*Synthesis of pGlu(OBn)$_{27}$-*b-*pSar$_{182}$*. The synthesis of pGlu(OBn)-*b*-pSar was adapted from the literature and modified. [1] 377.5 mg (1.43 mmol, 25 eq) Glu(OBn) NCA were weighed into a pre-dried Schlenk tube in a nitrogen counterstream and dried under high vacuum for 30 min. 3 mL dry DMF were added, and the solution was cooled to 0 °C. The reaction mixture was protected from light throughout the reaction time. 6.7 µL (0.06 mmol, 1 eq) neopentylamine in 1 mL DMF were added from a stock solution with a syringe and the reaction mixture was stirred at 0 °C with a slight stream of nitrogen attached. The progress of the polymerization was monitored by infrared (IR) spectroscopy. When the carbonyl stretching vibration bands of the monomer at 1853 $cm^{-1}$ and 1786 $cm^{-1}$ could not be detected anymore, 1188 mg (10.32 mmol, 200 eq) of the second monomer, sarcosine NCA, were added immediately, together with an additional 5 mL of dry DMF. The progress of the polymerization was monitored by IR spectroscopy. Directly after completion of the reaction, 50% of the polymer solution was precipitated into cold diethyl ether and centrifuged (4500 rpm, 4 °C, 15 min). After discarding the liquid fraction, diethyl ether was added again, and the polymer was resuspended in an ultrasonic bath. The suspension was centrifuged, resuspended, and centrifuged again. After the third centrifugation, the polymer was dissolved in Milli-Q water and lyophilized. The product was obtained as a colorless solid (457 mg, 89%). The other 50% of the reaction mixture was acetylated as described in the next paragraph.

*End group acetylation of pGlu(OBn)$_{27}$-b-pSar$_{182}$*. The procedure for the acetylation of terminal amine groups of polypept(o)ides was adapted from literature and modified.[2] When complete conversion of the last monomer was indicated by IR spectroscopy, 50% of the reaction mixture was precipitated in diethyl ether immediately, while 20 eq (80 µL, 0.57 mmol) triethylamine and 10 eq (27 µL, 0.29 mmol) acetic anhydride were added to the remaining 50%. This solution was stirred overnight at room temperature and afterwards precipitated in diethyl ether and centrifuged (4500 rpm, 4 °C, 15 min). After two resuspension and centrifugation steps, the polymer was dialyzed against Milli-Q water for three days (molecular weight cut-off (MWCO) 3.5 kDa) and lyophilized. The product was obtained as a colorless solid (453 mg, 89%).

$^1$H NMR (400 MHz, DMSO-$d_6$): $\delta$ [ppm] = 8.45-8.05 (s, br, 14H (1n), h), 7.35-7.15 (m, 134H (5n), a), 5.10-4.90 (m, 55H (2n), b), 4.50-3.80 (m, 391H (1n+2m), d, e), 3.00-2.65 (m, 557H (3m), f), 2.30-1.85 (m, 81H (4n), c), 0.84 (s, 9H, g).

GPC in HFIP (vs. PMMA standards) of pGlu(OBn)$_{27}$: $M_n$ = 9.6 kg/mol, Đ = 1.08.

GPC in HFIP (vs. PMMA standards) of pGlu(OBn)$_{27}$-*b*-pSar$_{182}$: $M_n$ = 34.5 kg/mol, Đ = 1.17.

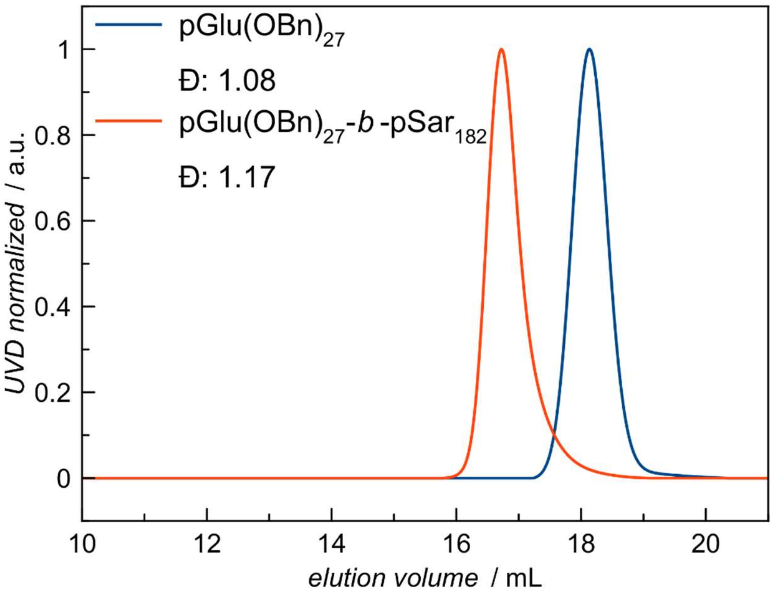


**Figure S1.** HFIP GPC elugrams (vs. PMMA standards) of pGlu(OBn)$_{27}$ and pGlu(OBn)$_{27}$-*b*-pSar$_{182}$.

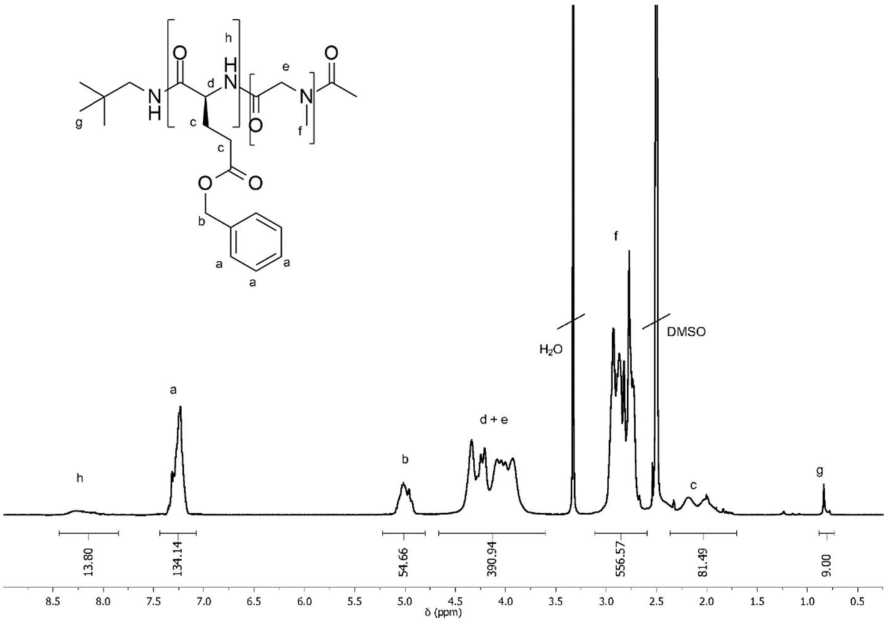


**Figure S2.** $^{1}$H NMR spectrum of pGlu(OBn)$_{27}$-*b*-pSar$_{182}$ in DMSO-$d_6$ measured at 400 MHz.

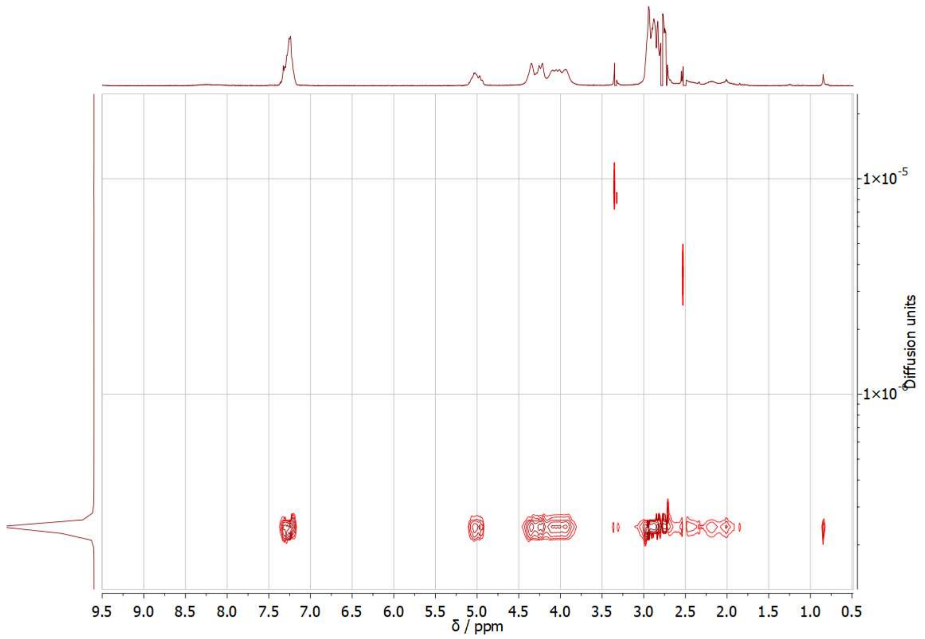


**Figure S3.** DOSY NMR spectrum of pGlu(OBn)$_{27}$-*b*-pSar$_{182}$ in DMSO-$d_6$ measured at 400 MHz.

*Fluorescent labeling of pGlu(OBn)$_{27}$*-b-*pSar$_{182}$ using the example of pGlu(OBn)$_{27}$*-b-*pSar$_{182}$-OG488*. The procedure for the labeling of block copolypept(o)ides with Oregon Green 488 succinimidyl ester (OG488-NHS) and Alexa Fluor 647 succinimidyl ester (AF647-NHS) was adapted from the literature and modified.[1] 29.7 mg (1.6 µmol, 1 eq) pGlu(OBn)$_{27}$-*b*-pSar$_{182}$ with free amine end group were weighed into a pre-dried Schlenk tube and dried under high vacuum for 2 h. Then, the polymer was dissolved in 1 mL of dry DMF. 0.3 µL (1.7 µmol, 1.1 eq) dry *N,N*-diisopropylethylamine (DIPEA) and subsequently 1.2 mg (2.3 µmol, 1.5 eq) OG488-NHS were added. The reaction mixture was stirred at room temperature under a nitrogen atmosphere for three days. The OG488-functionalized polymer was separated from free OG488 by column chromatography using sephadex LH-20 as stationary phase and DMSO as eluent and then dialyzed against Milli-Q water for three days (MWCO 3.5 kDa) and lyophilized (20.5 mg, 68% yield).

*Synthesis and modification of pGlu(OtBu)$_{32}$*-b-*pSar$_{202}$*. The syntheses were carried out analogously to the polymerization and post-polymerization reactions described above for *pGlu(OBn)$_{27}$*-b-*pSar$_{182}$*. During the polymerization of Glu(O*t*Bu) NCA, a mixture of DMF (67%) and THF (33%) was used as solvent. pGlu(O*t*Bu)$_{32}$-*b*-pSar$_{202}$ was obtained as a colorless solid (473 mg, 82% yield).

$^{1}$H NMR (400 MHz, $CDCl_3$): $\delta$ [ppm] = 8.30 (s, 24H (1n), g), 4.39 – 3.89 (m, 427H (1n+2m), c+d), 3.16 – 2.82 (m, 607H (3m), e), 2.25 (d, $J$ = 30.5 Hz, 296H (2n), b), 1.50 – 1.39 (m, 267H (9n), a), 0.96 – 0.87 (m, 9H, f).

GPC in HFIP (vs. PMMA standards) of pGlu(O*t*Bu)$_{32}$: $M_n$ = 7.7 kg/mol, Đ = 1.19.

GPC in HFIP (vs. PMMA standards) of pGlu(O*t*Bu)$_{32}$-*b*-pSar$_{202}$: $M_n$ = 34.2 kg/mol, Đ = 1.36.

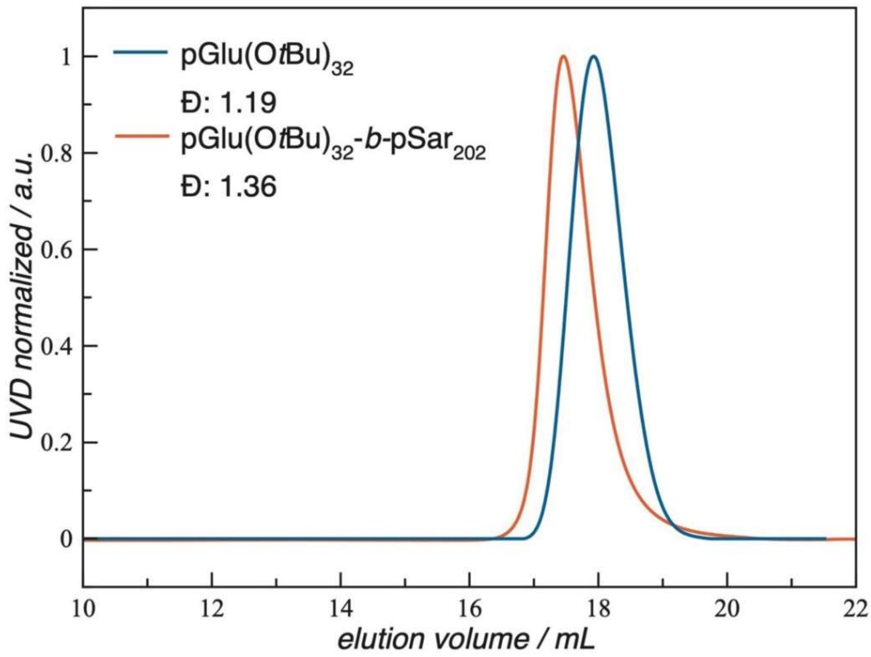


**Figure S4.** HFIP GPC elugrams (vs. PMMA standards) of pGlu(O*t*Bu)$_{32}$ and pGlu(O*t*Bu)$_{32}$-*b*-pSar$_{202}$.

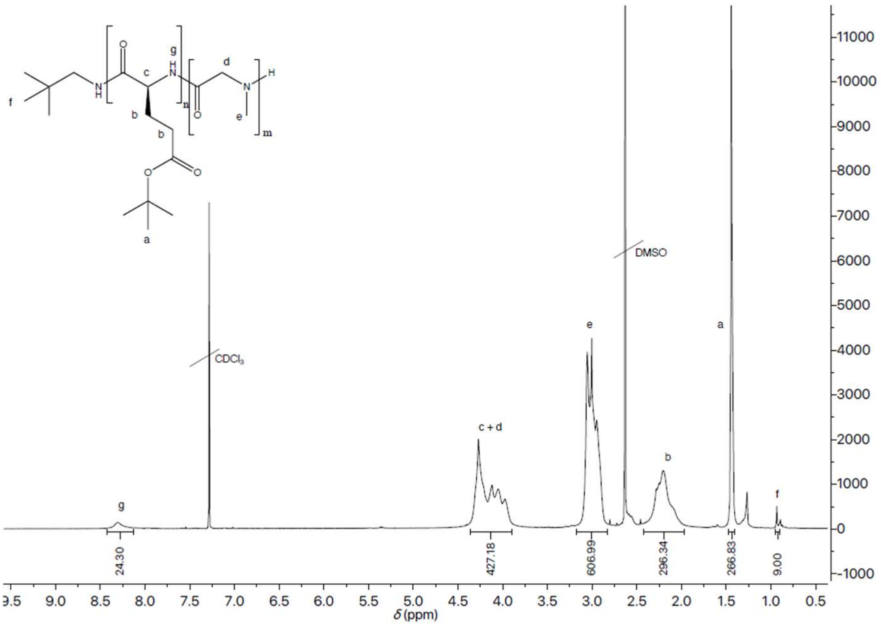


**Figure S5.** $^{1}$H NMR spectrum of pGlu(O*t*Bu)$_{32}$-*b*-pSar$_{202}$ in $CDCl_3$ measured at 400 MHz.

### 2. Determination of encapsulation efficiency and drug release profiles

#### The Drug Encapsulation Efficiency (EE %)

The encapsulation efficiency was calculated using the following equation:

$$\text{EE \%} = \frac{\mathrm{m}_{total} - \mathrm{m}_{free}}{\mathrm{m}_{total}} \times 100\ \%$$

Where $\mathrm{m}_{total}$ is the total amount of drug used in the synthesis and $\mathrm{m}_{free}$ is the amount of free drug.

For the quantification of different drugs, stock PM solutions at given drug concentrations were prepared and subjected to solid-phase extraction, separating free drug and encapsulated drug, and the fractions were analyzed by UPLC. That enabled a determination of the free drug concentration from the area under the curve (AUC), using the calibration plots below (in which the data allowed linear fitting). This concentration was then used to calculate the drug encapsulation efficiency.

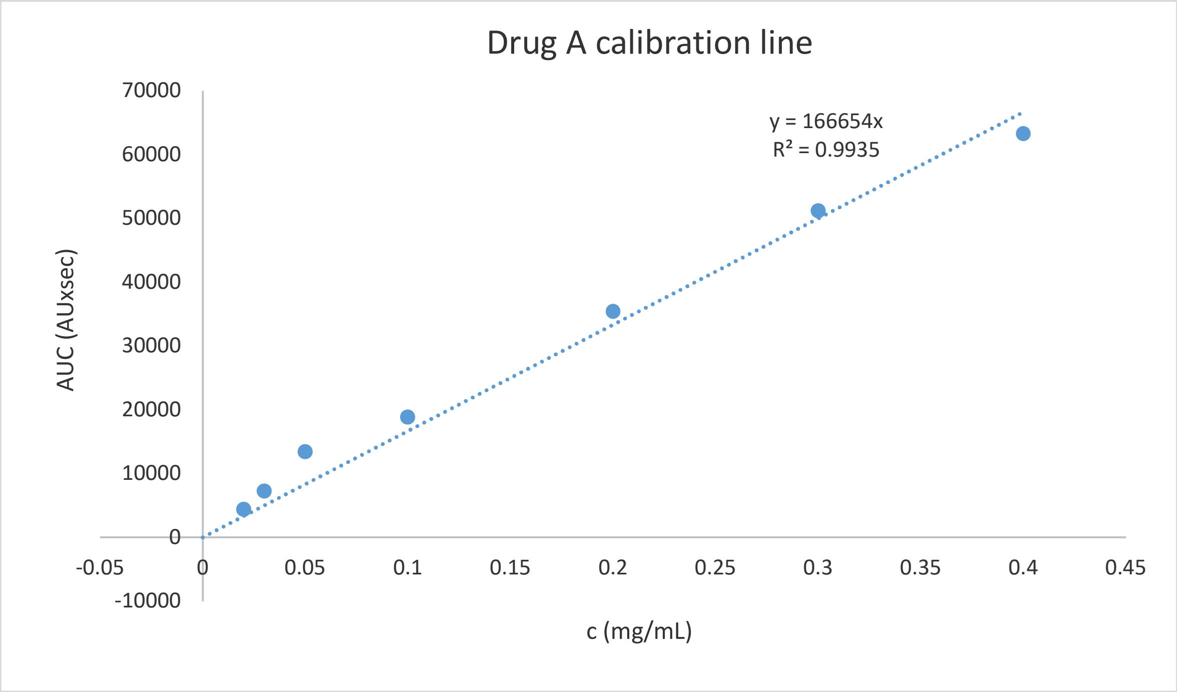


**Figure S6A:** UPLC standard calibration curve of Drug A.

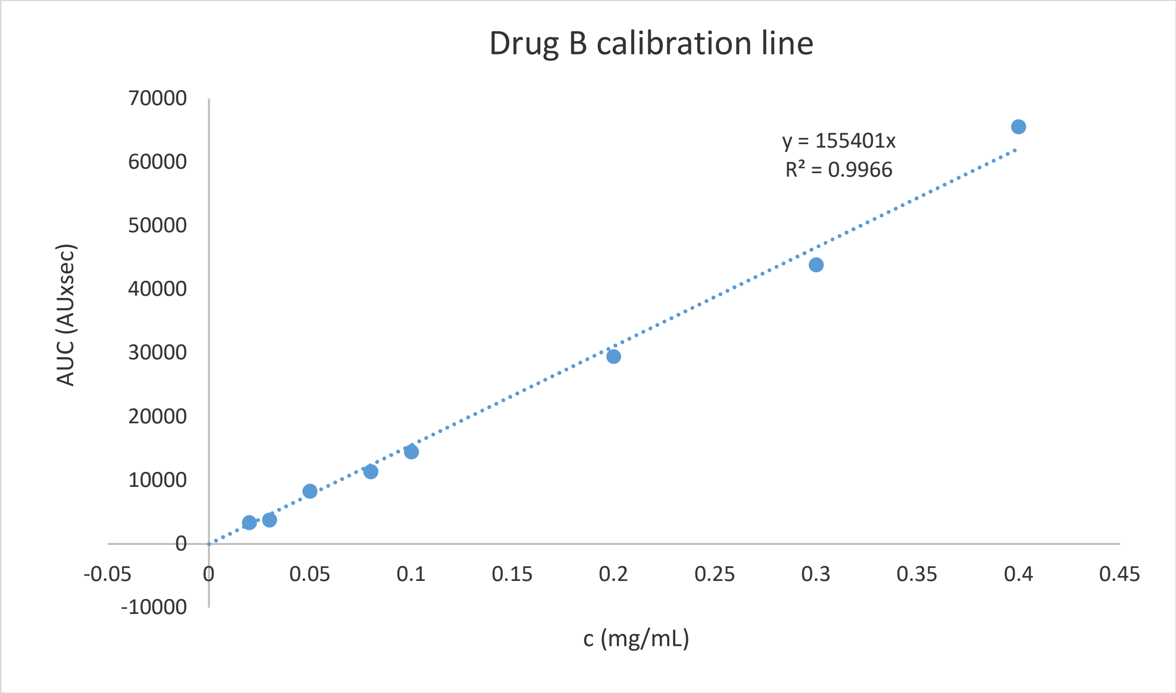


**Figure S6B:** UPLC standard calibration curve of Drug B.

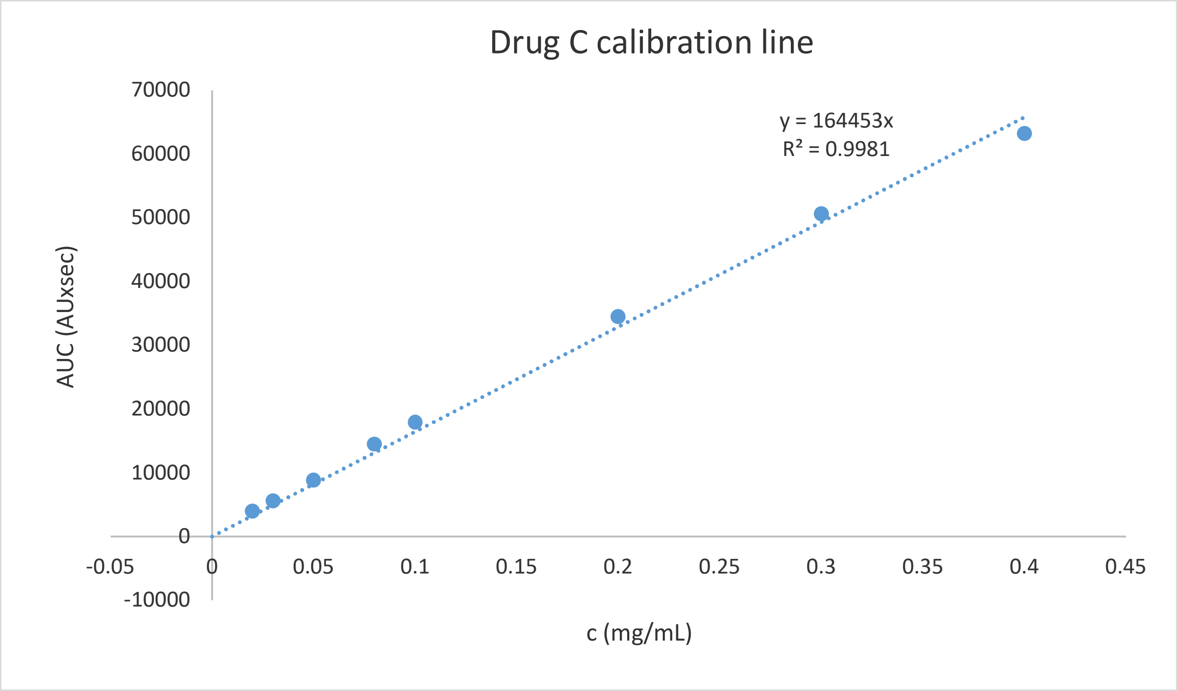


**Figure S6C:** UPLC standard calibration curve of Drug C.

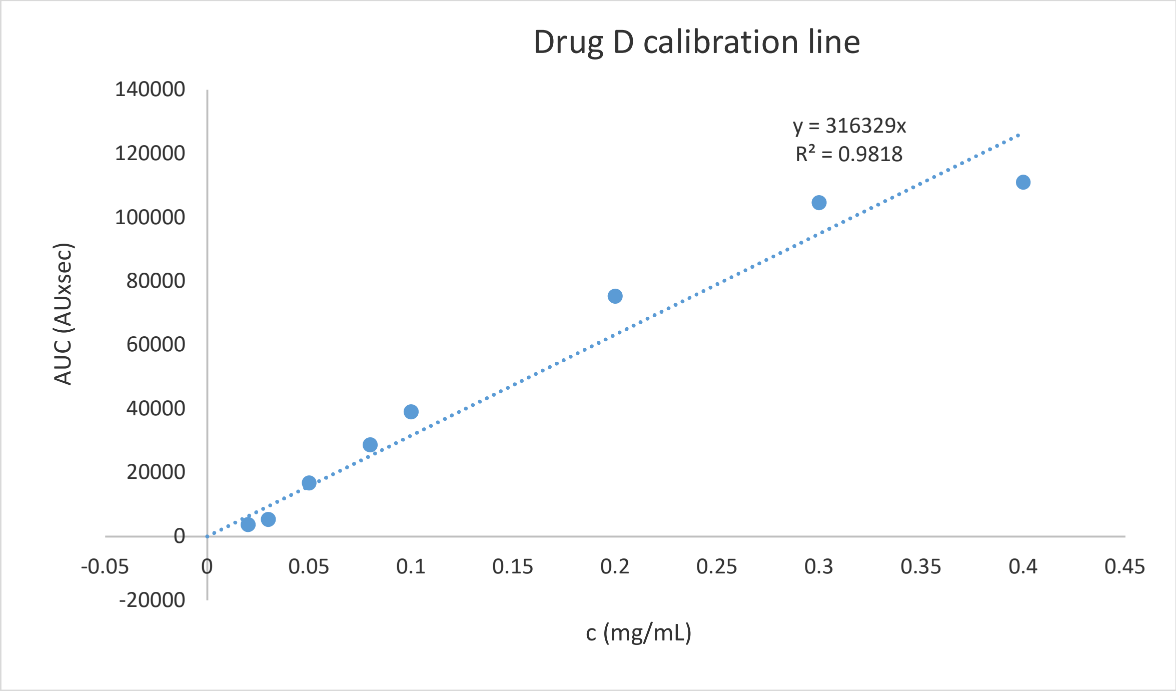


**Figure S6D:** UPLC standard calibration curve of Drug D.

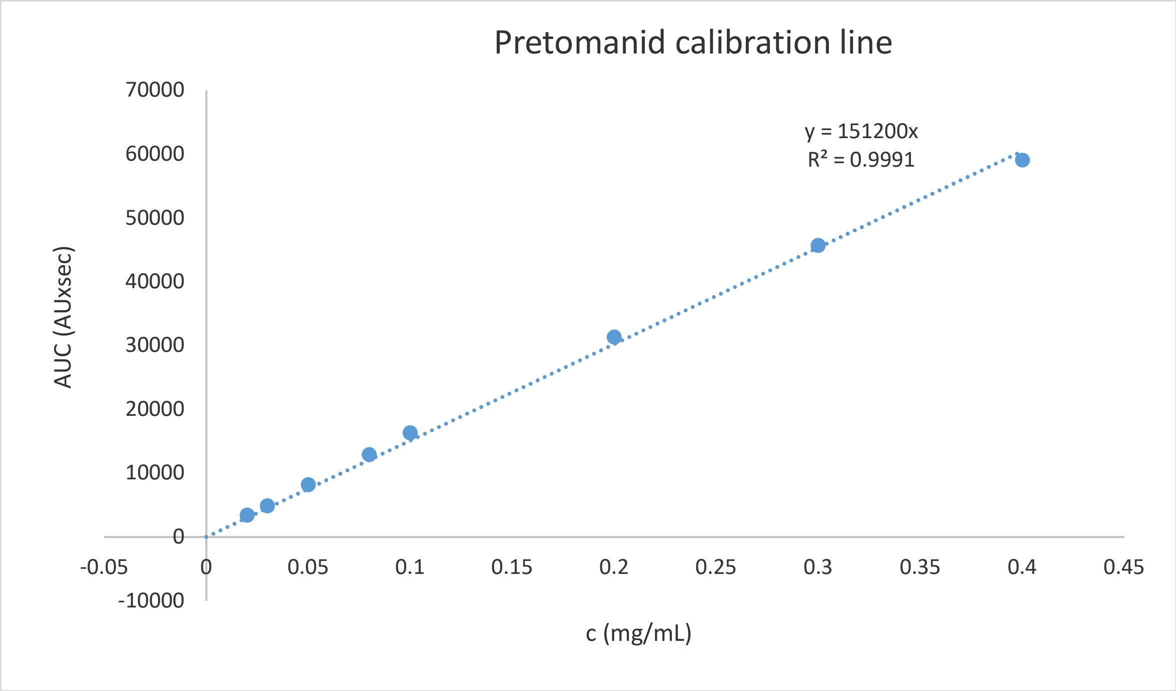


**Figure S6E:** UPLC standard calibration curve of Pretomanid.

### Drug Release from Micelles

The release of each drug (A, B, C, D and Pretomanid) from the PeptoMicelles was performed at 37 °C in PBS buffer. The micellar solution (200 μL, 25 mg/mL) was added into a Dialysis Membrane (MWCO: 3.5 kD) and stirred against 50 mL PBS. At different timepoints, the complete solution (50 mL) was removed and filtered, using SEP-PAK® Cartridge (C18) to get rid of PBS salt. The SEP units were flushed with acetonitrile (1 mL) to obtain the released drug, and each sample was further analyzed by UPLC, using the calibration curves determined before for each individual drug (see S6A-E).

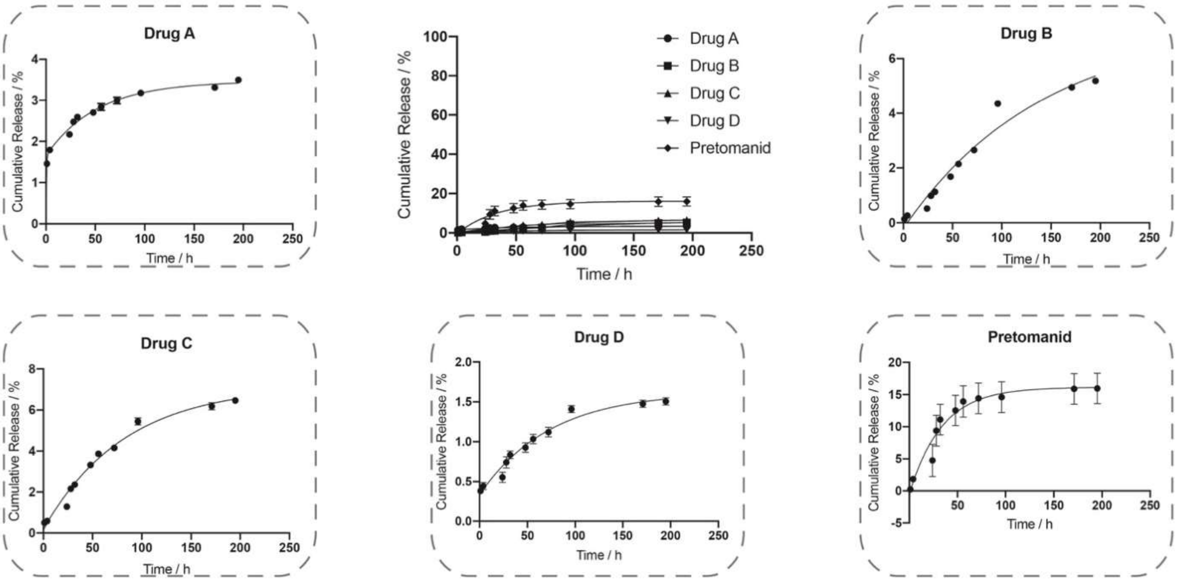


**Figure S7:** Drug release profiles of drug loaded PeptoMicelles. All experiments were conducted in triplicate.

### 3. Additional fluorescence cross-correlation spectroscopy (FCCS) data

Additional FCCS measurement data for PeptoMicelles formed from pGlu(OBn)$_{27}$-*b*-pSar$_{182}$ without drug-loading in water and with addition of 10 vol% DMSO, as well as for the micellar NP formulation of drug D in water, are shown below. For comparison, FCCS measurement data for micelles formed from pGlu(O*t*Bu)$_{32}$-*b*-pSar$_{202}$ in human blood plasma are shown, demonstrating that stability under these conditions is reduced when aromatic groups are missing. All samples were incubated in the respective medium for one week at room temperature prior to measurement.

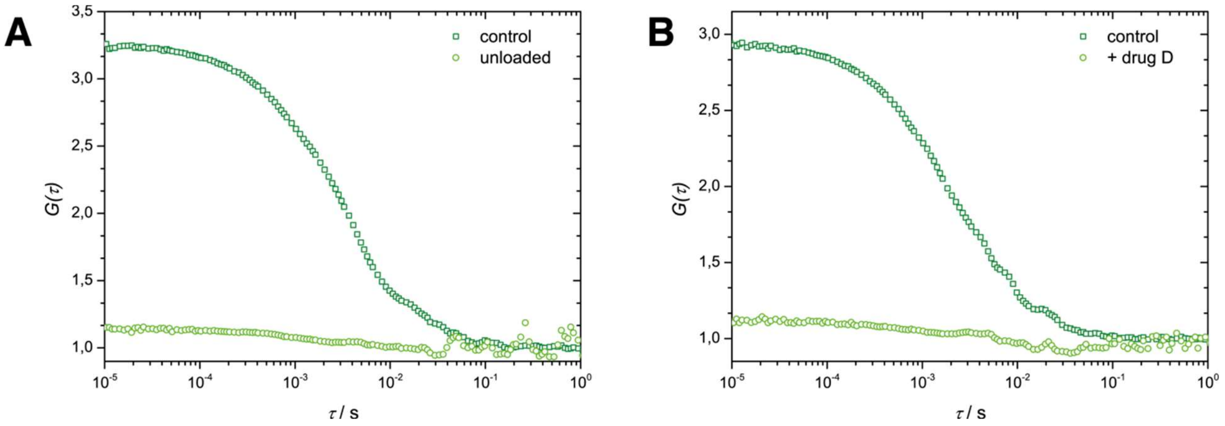


**Figure S8.** Cross-correlation functions (only) derived from FCCS measurements of non-loaded PeptoMicelles (**A**) and the micellar formulation of drug D (**B**) in water. Dark green (squares): Positive control measurements for a NP formulation that was labeled with both dyes after incubation in water at room temperature for one week. Light green (circles): Measurements after mixing of NP formulations labeled individually with OG488 or AF647 and incubation in water at room temperature for one week.

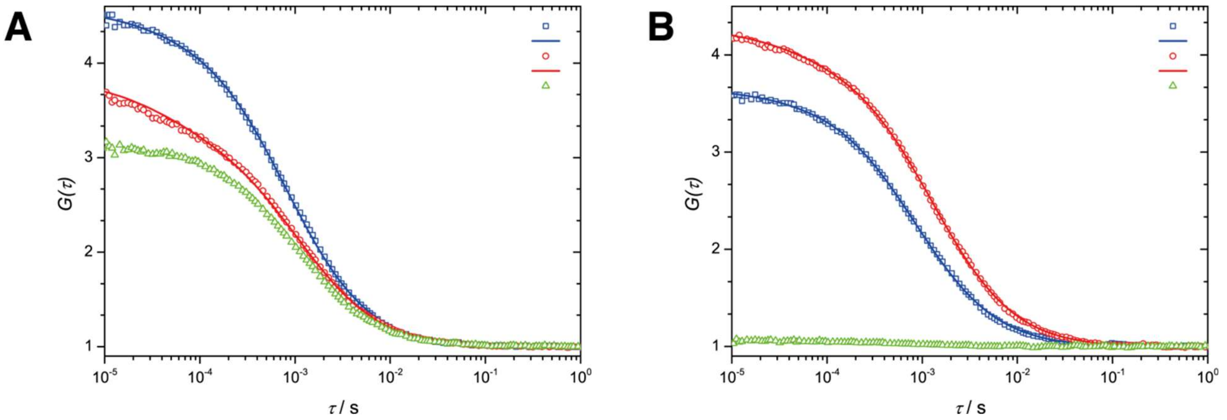


**Figure S9.** Auto- and cross-correlation functions derived from FCCS measurements of non-loaded PeptoMicelles in water with addition of 10 vol% DMSO. **A:** Positive control measurement for a NP formulation that was labeled with both dyes after incubation in 10% DMSO/water at room temperature for one week. **B:** Measurement after mixing of NP formulations labeled individually with OG488 or AF647 and incubation in 10% DMSO/water at room temperature for one week. Blue: Oregon Green 488 (OG488) experiment (squares) and fit function (line). Red: Alexa Fluor 647 (AF647) experiment (squares) and fit function (line). Green: Cross-correlation of both fluorescence signals.

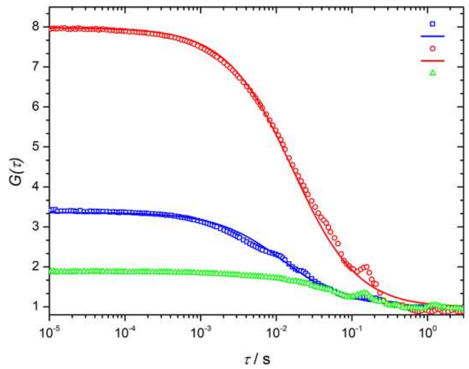


**Figure S10.** Auto- and cross-correlation functions derived from FCCS measurements in human blood plasma of NP formulations formed from pGlu(O*t*Bu)$_{32}$-*b*-pSar$_{202}$ (no drug load, samples containing each fluorophore were individually prepared and subsequently mixed and incubated in human blood plasma). Blue: Oregon Green 488 (OG488) experiment (squares) and fit function (line). Red: Alexa Fluor 647 (AF647) experiment (squares) and fit function (line). Green: Cross-correlation of both fluorescence signals.

### 4. Improved synthetic route for the larger-scale synthesis of pretomanid analogue D

The reported synthesis[3] of analogue D involved sequential Sonogashira coupling of bromide **1** with ethynyl(trimethylsilane), desilylation (TBAF), and Sonogashira coupling of the resulting terminal acetylene intermediate with the appropriately-substituted iodobenzene. The improved larger scale route reduced these three steps into one, *via* Sonogashira coupling of bromide **1** with 1-ethynyl-4-(trifluoromethoxy)benzene, giving analogue D in 87% yield.

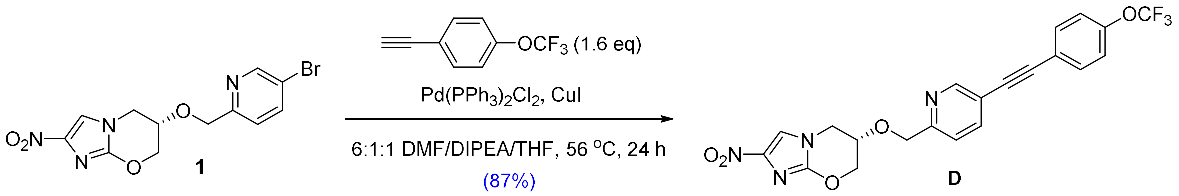


**(6*S*)-2-Nitro-6-[(5-{[4-(trifluoromethoxy)phenyl]ethynyl}pyridin-2-yl)methoxy]-6,7-dihydro-5*H*-imidazo[2,1-*b*][1,3]oxazine (D)**. A stirred mixture of (6*S*)-6-[(5-bromopyridin-2-yl)methoxy]-2-nitro-6,7-dihydro-5*H*-imidazo[2,1-*b*][1,3]oxazine[4] (**1**) (2.004 g, 5.64 mmol), $Pd(PPh_3)_2Cl_2$ (273 mg, 0.389 mmol) and copper(I) iodide (109 mg, 0.572 mmol) in anhydrous DMF (40 mL), anhydrous distilled THF (10 mL) and *N*,*N*-diisopropylethylamine (6.1 mL, 35 mmol) was degassed with a vacuum pump for 20 min and then sealed under $N_2$. 1-Ethynyl-4-(trifluoromethoxy)benzene (1.40 mL, 9.14 mmol) was added by syringe and the sealed mixture was stirred at 56 °C for 24 h, then cooled, added to ice/aqueous $NaHCO_3$ (250 mL), and extracted with $CH_2Cl_2$ (2 x 200 mL then 3 x 100 mL). The combined extracts were evaporated to dryness under reduced pressure (30 °C) and the residue was chromatographed on silica gel. Elution with

10-75% EtOAc/petroleum ether first gave foreruns, and then further elution with EtOAc gave the crude product (2.61 g), which was chromatographed again twice on silica gel. Elution with $CH_2Cl_2$ first gave foreruns, and then further elution with 2% MeOH/$CH_2Cl_2$ gave the product, which was recrystallized three times from MeOH/$CH_2Cl_2$/hexane to give **D** (2.27 g, 87%) as a pale yellow solid.

mp: 211-214 °C.

$^1$H NMR [$(CD_3)_2SO$]: $\delta$ [ppm] = 8.72 (d, $J$ = 1.4 Hz, 1 H), 8.05 (s, 1 H), 8.00 (dd, $J$ = 8.1, 2.2 Hz, 1 H), 7.73 (d, $J$ = 8.9 Hz, 2 H), 7.48-7.41 (m, 3 H), 4.81 (d, $J$ = 13.7 Hz, 1 H), 4.77 (d, $J$ = 13.8 Hz, 1 H), 4.72 (dt, $J$ = 12.0, 2.5 Hz, 1 H), 4.50 (br d, $J$ = 11.8 Hz, 1 H), 4.38-4.31 (m, 2 H), 4.26 (dd, $J$ = 13.7, 3.5 Hz, 1 H); HPLC purity: 97.9%.

### 5. Additional zebrafish (ZF) data

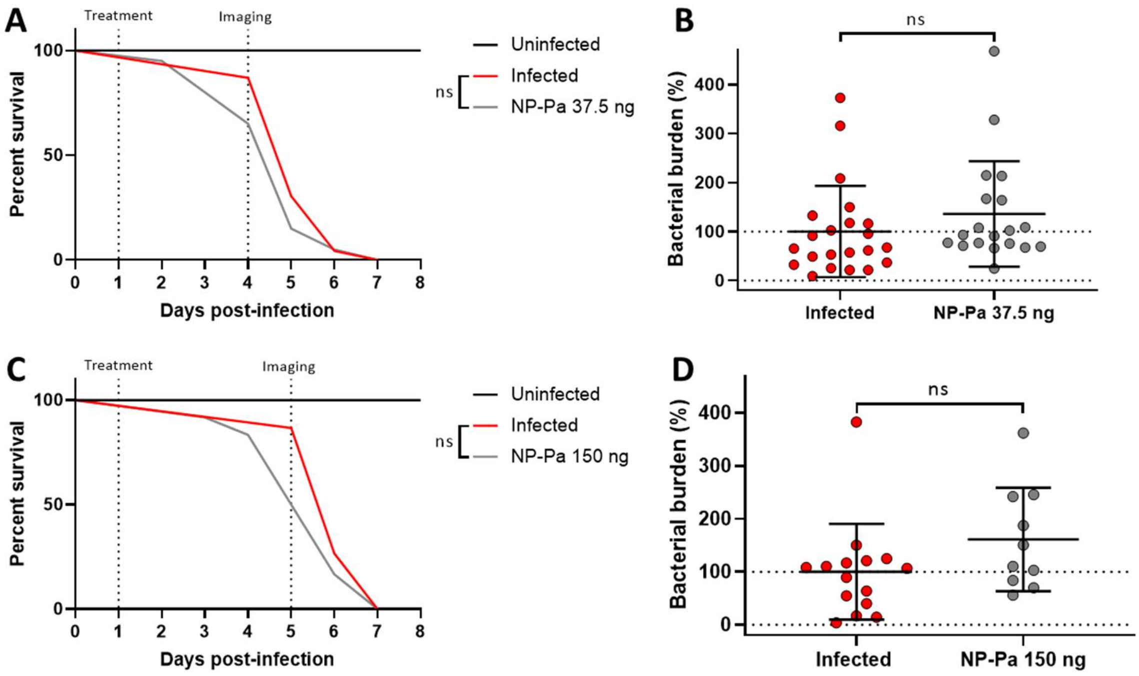


**Figure S11.** *In vivo* activity study of pretomanid (Pa), as a micellar drug formulation, against *Mm* in the zebrafish embryo model. Fish infected by the blood (PCV). **A:** Survival analysis for the low dose (37.5 ng) tested. N (zebrafish per group at day of infection) ≥ 20. **B:** FPC analysis of bacterial burden at day 4 post-infection for the low dose tested. N (zebrafish per group at day of imaging) ≥ 19. Statistics; Mann-Whitney test (non-parametric t test). **C:** Survival analysis for the high dose (150 ng) tested. N (zebrafish per group at day of infection) ≥ 12. **D:** FPC analysis of bacterial burden at day 5 post-infection for the low dose tested. N (zebrafish per group at day of imaging) ≥ 10. Statistics; Mann-Whitney test (nonparametric t test).

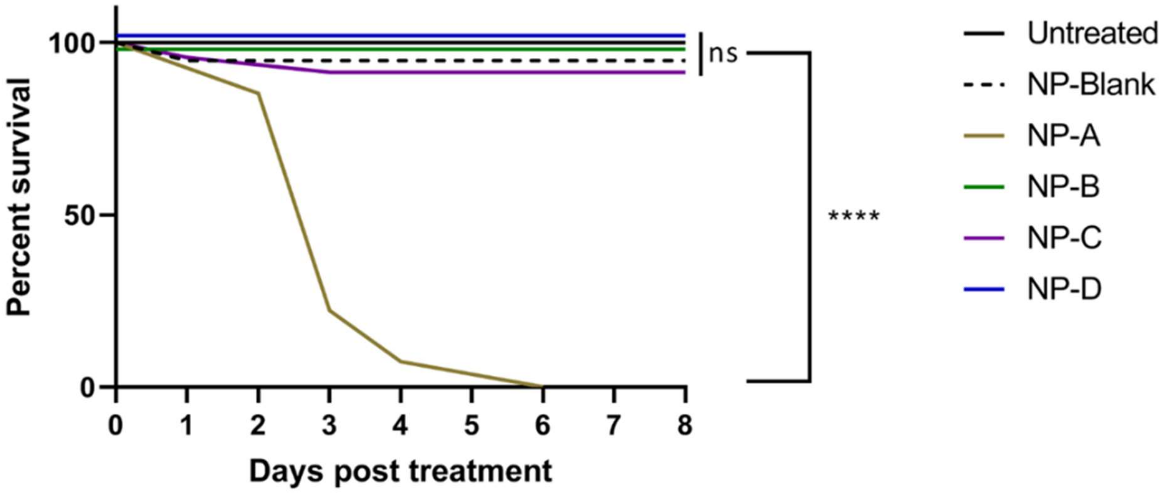


**Figure S12.** Toxicity/tolerability screen survival analysis for the four pretomanid derivatives as micellar NP formulations in healthy uninfected zebrafish larvae. The dosage in each case was equivalent to 75 ng of drug, administered on day two post-fertilization; blank NP (micelles without drug) were used as a control. N (zebrafish per group) ≥ 19.

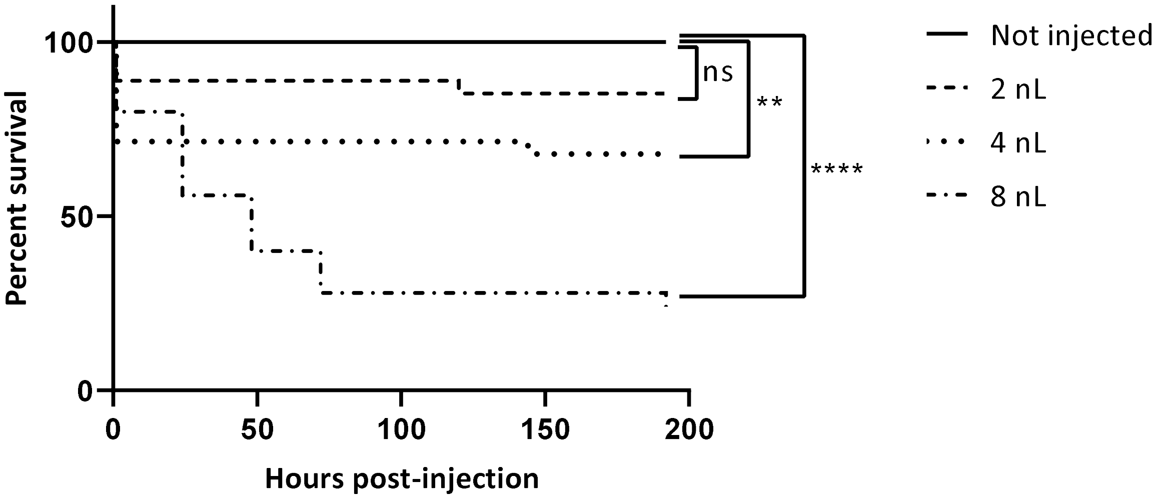


**Figure S13.** Vehicle tolerability study of PEG400:DMSO (4:1 ratio) in uninfected zebrafish embryos. Solution injected by the blood (PCV) at 48 h post-fertilization. The group injected with the lowest volume, 2 nL, was the only group not significantly different from the non-injected control group. Therefore, 2 nL was considered the maximum tolerated volume of this solution for injections. Consequently, the free form of drugs A and D was solubilized in 4:1 PEG400:DMSO and no more than 2 nL was administered, regardless of the drug concentration. N (zebrafish per group) ≥ 12.

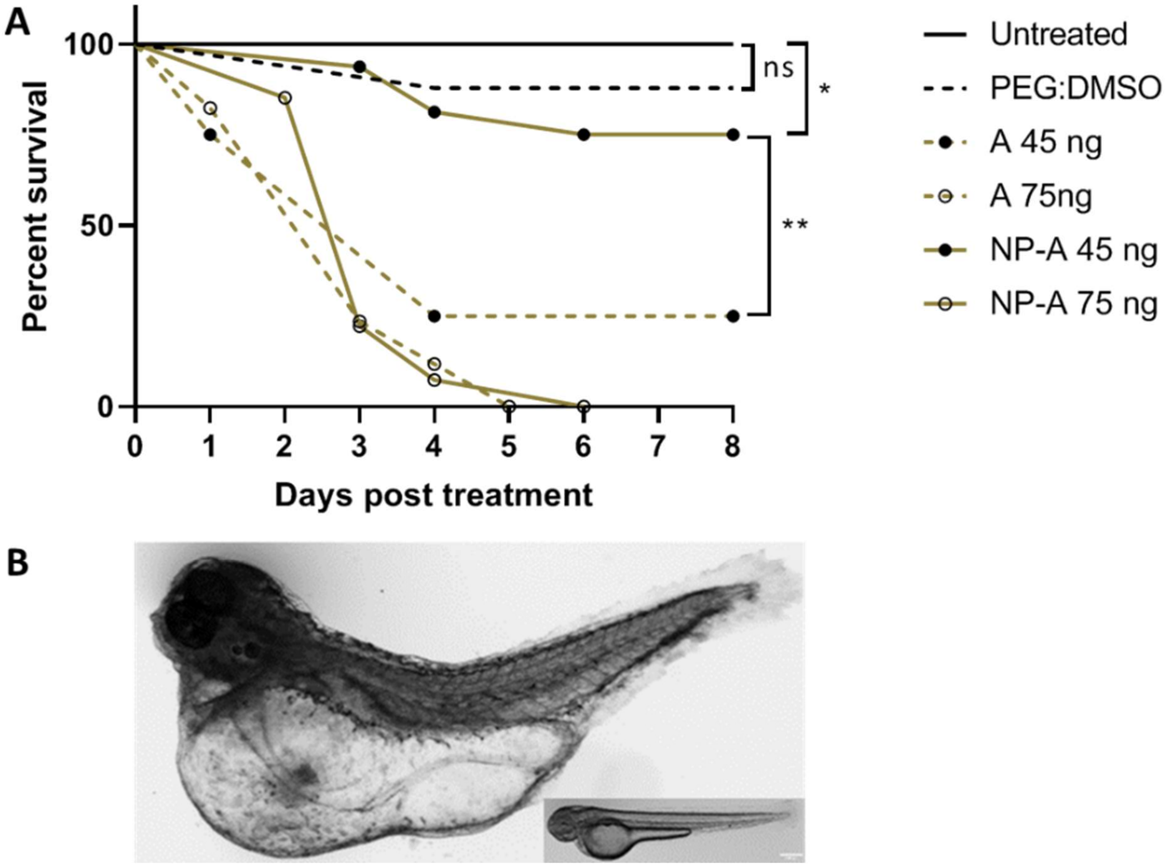


**Figure S14.** Toxicity/tolerability screen of drug A, free *vs.* micellar drug formulation. **A:** Survival analysis. Drug A administered at two doses (75 ng and 45 ng) in both formulations. The micellar formulation significantly reduced the mortality at 45 ng compared to the equivalent dose in free form (PEG400:DMSO as solvent), but not at 75 ng. The vehicle control group for the free drug was not significantly different from the untreated control group, vehicle control and free drug groups had a total injection volume of 2 nL. N (zebrafish per group) ≥ 12. **B:** Zebrafish injected with 40 ng of drug A in the micellar formulation. The image is acquired eight days post PCV infection with dsRed *Mm* (seven days after treatment) and not directly related to the survival curve in **A**. The image clearly illustrates the edemas associated with exposure to drug A. The inset, as a comparison, shows a normal healthy embryo with no signs of toxicity.

### 6. Pharmacokinetic studies in mice

Pharmacokinetic studies were performed at Wuxi AppTec Co., Ltd., Cranbury, New Jersey. All procedures relating to animal handling, care, and treatment were performed according to the guidelines approved by the Institutional Animal Care and Use Committee of Wuxi AppTec Co. following the guidance of the Association for Assessment and Accreditation of Laboratory Animal Care.

Two formulations of drug D were studied using groups of three female 6-8 weeks old C3HeB/FeJ mice (sourced from The Jackson Laboratory). The test compound was first encapsulated in polymeric micelles and supplied as an aqueous solution (comprising 25 mg/mL nanoparticles, with 30% drug encapsulation, i.e., 7.5 mg/mL of drug), ready for intravenous dosing at 40 mg/kg. The free drug was formulated as a 20 mg/mL solution in 4:1 (v/v) PEG-400/DMSO, also for intravenous dosing at 40 mg/kg. Blood samples were collected at 0.083, 0.5, 1, 2, 4, 8, 24, and 48 h, transferred into prechilled K2-EDTA tubes, and kept on ice until being processed for plasma by centrifugation at 4 °C. Plasma samples were stored at -70 °C prior to analysis by LC-MS/MS, and the PK parameters were determined using Phoenix WinNonlin software (version 6.3) based on noncompartmental analysis.

**Table S1.** Mouse pharmacokinetic parameters for drug D formulations following a single intravenous dose.

| Form | $C_{max}$ (μg/mL) | $t_{1/2}$ (h) | $AUC_{inf}$ (μg·h/mL) |
|---|---|---|---|
| free D | 16.3 | 48 | 776 |
| NP-D | 18.8 | 55 | 890 |

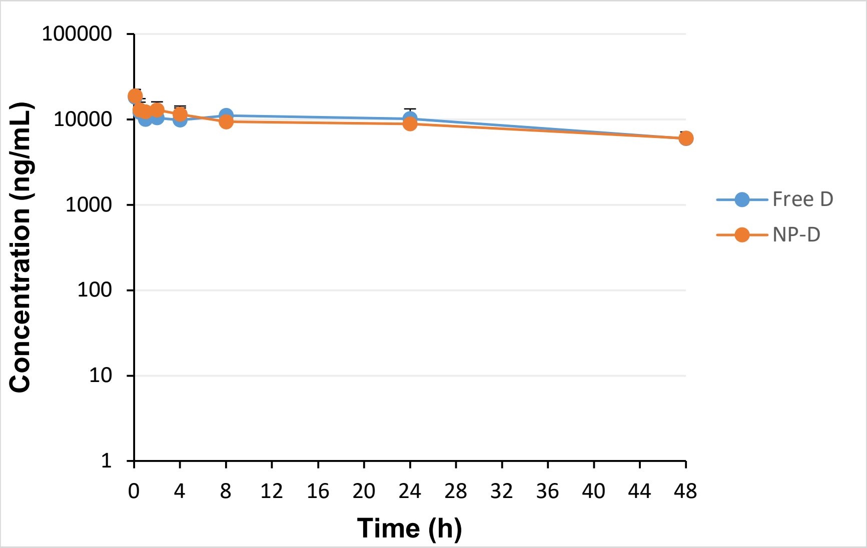


**Figure S15.** Plasma concentration-time profiles for drug D formulated in 4:1 PEG400/DMSO (free) or in polymeric micelles following intravenous injection at 40 mg/kg in C3HeB/FeJ mice.

### 7. Additional mouse data

Relative area of inflammation analysis, haematoxylin, and Eosin staining, followed by outlining region of interest (ROI) for healthy and inflamed tissue, and histological analysis (Haematoxylin and Eosin, "oil red o" and Ziehl–Neelsen stain) of lung sections from one of seven mice in the infected control group, derived from the second experiment.

*H&E Staining.* Cryo sections were rehydrated by dipping in Aqua dest (A. dest), the sections were coloured for 10 min using Hemalaun solution, and then washed in tap water, which led to blue staining of the nuclei. Sections were rinsed briefly in A. dest, counterstained with 1% Eosin for 1 min showing cell plasma in red, passed through A. dest, and dehydrated in ascending concentrations of alcohol (dipping twice 2 min each in 96% EtOH and 100% EtOH) and Xylene. Finally, the sections were embedded in Entellan and observed under the light microscope BX41 (Olympus, Tokyo, JP).

*Staining of lipid droplets (or bodies) in lung tissue.* Lipid staining of lung sections obtained from infected treated and non-treated mice was performed using "oil red o" at day 44 post-infection (nuclei- blue, lipid bodies- red). "Oil red o" (ORO) staining was used to identify triglycerides, triacylglycerols and lipid droplets in cryosections. Sections were rinsed in PBS and treated with freshly prepared ORO working solution (0.5% (w/v) in 60% triethylphosphate), then diluted to 0.35 (v/v) ORO in A. dest for 30 min. Sections were washed three times in A. dest, stained with Mayer´s Haematoxylin for 10 min, and briefly rinsed in A. dest. Finally, slides were mounted in aq. Kaiser´s glycerol gelatin in advance of microscopy using a BX41 light microscope (Olympus, Tokyo, JP).

*Ziehl-Neelsen (ZN) stain.* ZN staining was used to identify *Mycobacterium* species in cryo sections. Rehydrated sections were flooded with carbol-fuchsin and passed over a heat source until the appearance of fumes. After 5 min of cooling, sections were washed in A. dest and flooded with 0.5% (v/v) HCl in 70% (v/v) Ethanol until the sections appeared to be decolorized to pale pink. After brief rinsing in A. dest, sections were treated with Löffler´s methylene blue solution for 1 min and rinsed again with A. dest, dipped briefly in 96% EtOH and Isopropanol, before washing with Xylol. Finally, dried sections were embedded in Entellan and examined using a BX41 light microscope (Olympus, Tokyo, JP).

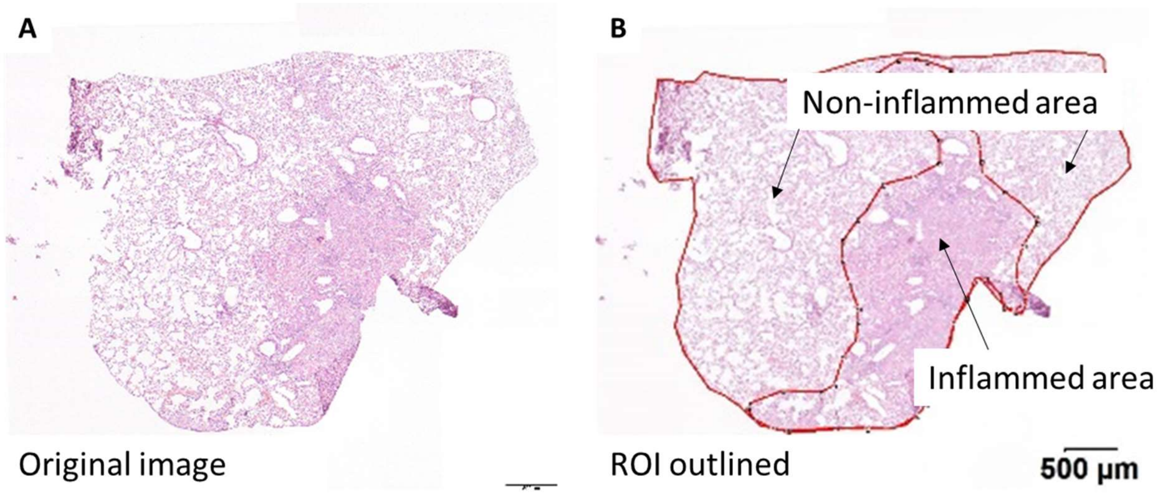


**Figure S16.** Example image of the relative area of inflammation analysis of lung sections from one of seven mice in the infected control group, derived from the second experiment. Hematoxylin and Eosin staining (**A**), followed by outlining region of interest (ROI) for healthy and inflamed tissue (**B**).

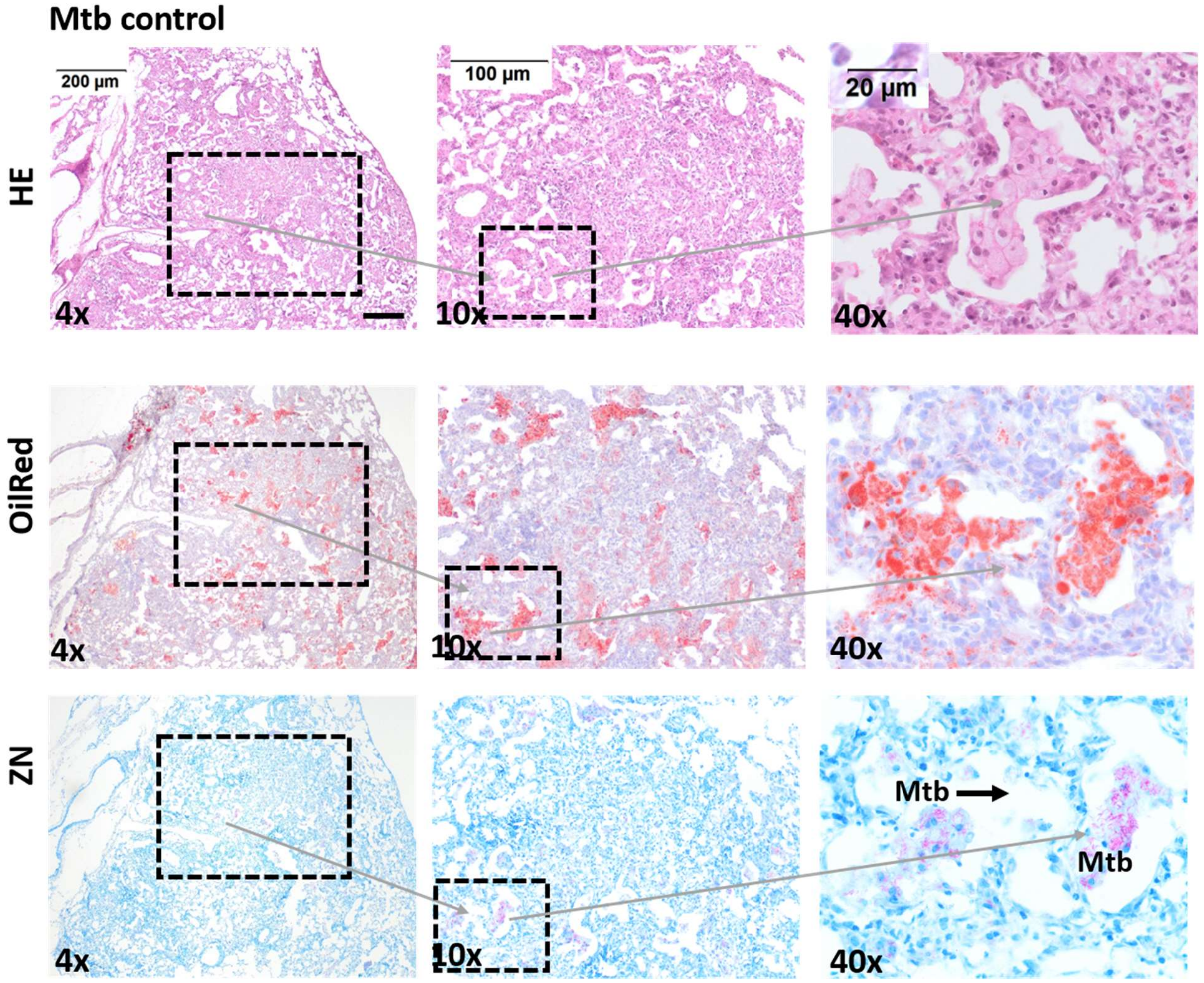


**Figure S17.** Histological analysis (Hematoxylin and Eosin, "oil red o" and Ziehl–Neelsen stain) of lung sections from one of seven mice in the infected control group, derived from the second experiment.

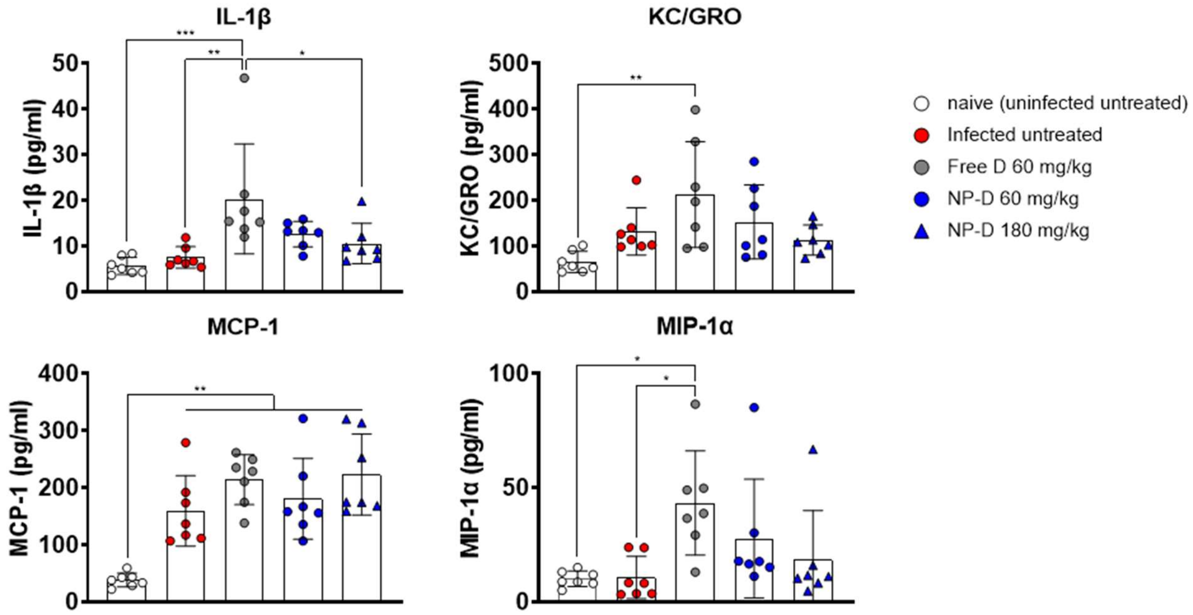


**Figure S18.** Inflammation data derived from lung samples for the second experiment. Concentrations of different inflammation markers in serum of mice on day 44 after *Mtb* H37Rv aerosol infection. The data given represent the mean and standard deviation from pooling two independent replicates per mouse (n = 7 mice per treatment group). Statistical analyses were performed by ordinary one-way ANOVA with Tukey's multiple comparisons test (The mean of each group is compared with the mean of every other group, only significant differences indicated in the graph).
